# Modality-specific circuits for skylight orientation in the fly visual system

**DOI:** 10.1101/638171

**Authors:** Gizem Sancer, Emil Kind, Haritz Plazaola, Jana Balke, Tuyen Pham, Amr Hasan, Lucas Münch, Thomas F. Mathejczyk, Mathias F. Wernet

## Abstract

In the fly optic lobe ∼800 highly stereotypical columnar microcircuits are arranged retinotopically to process visual information. Differences in cellular composition and synaptic connectivity within functionally specialized columns remains largely unknown. Here we describe the cellular and synaptic architecture in medulla columns located downstream of photoreceptors in the ‘dorsal rim area’ (DRA), where linearly polarized skylight is detected for guiding orientation responses. We show that only in DRA medulla columns, both R7 and R8 photoreceptors target to the bona fide R7 target layer where they form connections with previously uncharacterized, modality-specific Dm neurons: Two morphologically distinct DRA-specific cell types (termed Dm-DRA1 and Dm-DRA2) stratify in separate sublayers and exclusively contact polarization-sensitive DRA inputs, while avoiding overlaps with color-sensitive Dm8 cells. Using the activity-dependent GRASP and trans-Tango techniques, we confirm that DRA R7 cells are synaptically connected to Dm-DRA1, whereas DRA R8 form synapses with Dm-DRA2. Finally, using live imaging of ingrowing pupal photoreceptor axons, we show that DRA R7 and R8 termini reach layer M6 sequentially, thus separating the establishment of different synaptic connectivity in time. We propose that a duplication of R7→Dm circuitry in DRA ommatidia serves as an ideal adaptation for detecting linearly polarized skylight using orthogonal e-vector analyzers.

## Introduction

The brain contains many different neuronal circuits specialized for computing discrete features that the sensory system extracts from the environment. Identifying the cellular elements forming these circuits and understanding how their synaptic connections enable these circuits to perform their specific computational function remains a major challenge ^1^. Both in mammals and invertebrate models, the visual system has long served as a powerful system for investigating the neuronal basis of specific computations covering different aspects of visual perception, like color vision ^2–4^, the detection of looming ^5–7^ or moving stimuli ^8^. In recent years, the quickly growing arsenal of molecular-genetic tools available in *Drosophila* has been used to dissect the neuronal circuits underlying specific visual behaviors via the cell type-specific visualization and/or manipulation of neuronal activity. Additionally, new tools for the detailed characterization of neuronal morphology ^9–11^ and synaptic connectivity ^12, 13^ have been developed. As a result, the computation of visual motion within the optic lobes downstream of the fly’s stereotypical unit eyes (ommatidia) has been dissected with great success, at a cellular and synaptic level ^8, 14^.

In the fly retina, expression of four different *Drosophila* Rhodopsins in the inner photoreceptors R7 and R8 leads to the formation of a retinal mosaic for color vision where two ommatidial subtypes with different spectral sensitivities (called ‘pale’ and ‘yellow’) are randomly distributed ^15^. Importantly, each point in space perceived by fly photoreceptors is represented by a matching columnar element within the neuropils of the optic lobes (lamina, medulla, and lobula complex), resulting in a retinotopic representation of the visual world ^16, 17^. The medulla alone contains more than 80 distinct cell types and represents the most complex part of the fly visual system ^18, 19^. The wiring diagram of medulla columns has been shown to be highly stereotypical, harboring a fixed set of both columnar and multicolumnar cell types ^20^. Only few light microscopic studies have revealed cell types that exist specifically in those columns post-synaptic to either pale or yellow ommatidia ^21, 22^, yet their computational role remains largely unknown. More importantly, nothing is known about differences in circuit architecture (connectivity, synaptic distribution) between pale and yellow medulla columns and how these affect their functional role in color vision ^23^.

In addition to stochastically distributed ommatidia, virtually all insect retinas contain a third, morphologically specialized subtype along the dorsal eye margin, the so-called ‘dorsal rim area’ (DRA) ^24–26^. In *Drosophila* DRA ommatidia, photoreceptors R7 and R8 are monochromatic, expressing the same UV Rhodopsin (Rh3) ^27^ and are polarization-sensitive due to untwisted light-sensitive structures (rhabdomeres) that serve as orthogonal analyzers (R7 vs R8) for detecting linearly polarized skylight (Figure 1A) ^28, 29^. Furthermore, *Drosophila* DRA ommatidia are both necessary and sufficient to mediate specific orientation behavior in response to polarized light, which serves as a navigational help during walking ^28^. Importantly, a sharp boundary exists between polarization-sensitive DRA ommatidia and color-sensitive non-DRA ommatidia (Figure 1B) ^24, 26, 30^. This strict division is already detectable during mid-pupal stages, where DRA inner photoreceptors express the transcription factor Homothorax (Hth) that, when over-expressed, is sufficient to transform the entire retinal mosaic into a homogeneous field of DRA ommatidia (Figure 1C) ^26, 31, 34^. Interestingly, R8 cells in DRA ommatidia (here referred to as DRA.R8) resemble R7 cells in that they express an R7 Rhodopsin (Rh3), and down-regulate the crucial R8-specific transcription factor Senseless (Sens) around mid-pupation ^26, 31^. Furthermore, only in the DRA, R8 axons target to the deeper medulla layer M6, known to be the R7 target layer across the medulla ^32^, whereas non-DRA R8 always terminate in layer M3 ^19^. Differences in layer targeting are known to be crucial for correct synaptic partner choice by photoreceptors ^33^. However, nothing is known about differences in synaptic partner choice between DRA and non-DRA R7 or R8 photoreceptor cells.

**Figure 1:**
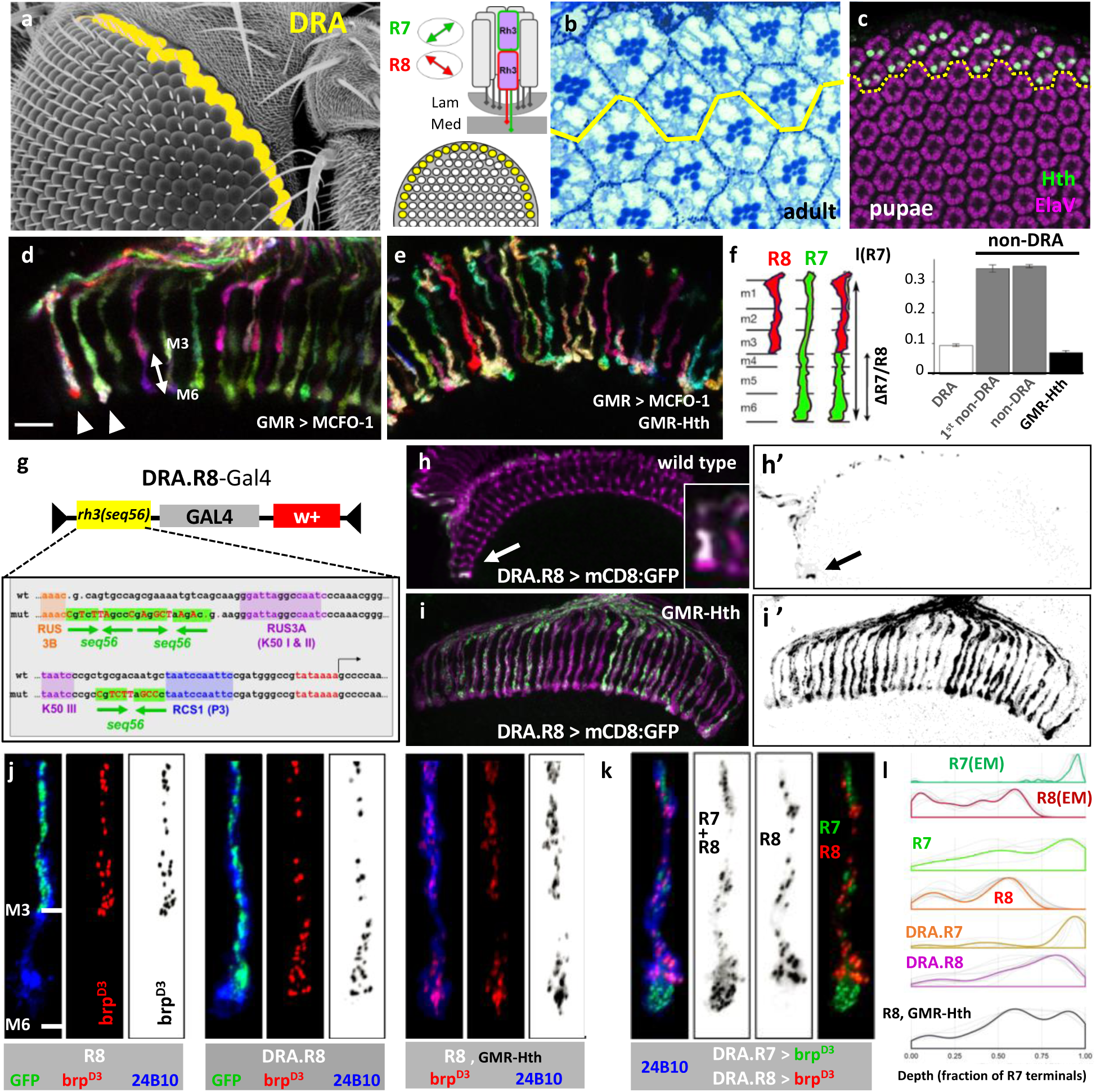
Presynaptic properties of DRA inner photoreceptors R7 and R8. **A.** Left: Electron micrograph depicting the approximate position of *Drosophila* polarization-sensitive DRA ommatidia (yellow). Right: Schematic depiction of one DRA ommatidium. Both inner photoreceptor cells R7 (green) and R8 (red) project axons into the medulla neuropil and contain the UV-Rhodopsin Rh3, usually found in ∼35% of non-DRA R7 cells. Double headed arrows indicate orthogonal arrangement of untwisted rhabdomeres in polarization-sensitive DRA.R7 versus DRA.R8 cells. **B.** Plastic section revealing the sharp boundary (yellow line) between DRA ommatidia (North) and non-DRA ommatidia (South); reproduced with permission from ^26^. Note the larger inner photoreceptor cell rhabdomere diameter inside the DRA. **C.** Whole-mounted pupal retina (50h APF) stained with Anti-ElaV (purple) and Anti-Homothorax (Hth, green) revealing a sharp boundary between DRA ommatidia (R7 and R8 labeled with Hth) and the rest of the developing eye (dorsal up). **D.** Adult whole-mounted brain with MCFO clones using photoreceptor-specific GMR-Gal4 revealing differences in R8 layer-specific targeting between non-DRA (target: layer M3) and DRA R8 cells (arrowheads, target: M6), resulting in a sharp DRA boundary in the medulla. **E.** Transformation of all ommatidia into the DRA fate using LongGMR-GFP:Hth (referred to as GMR-Hth) resulting in all R8 cells targeting to layer M6. **F.** Quantification of layer targeting. Left: Distance between R7 and R8 terminals (ΔR7/R8) is divided by the total length of R7 in the medulla l(R7). Right: plot depicting the sharp boundary between R8 targeting in the DRA, first non-DRA columns, random non-DRA columns, and GMR-Hth flies (indistinguishable from DRA). **G.** Description of mutant *rh3* promoter generated to specifically target Gal4 expression to DRA.R8 cells (DRA.R8-Gal4): Three palindromic R7 repressor sites (Pros binding sites) from *rh5* and *rh6* promoters (seq56) were introduced (gift from Dmitri Papatsenko, unpublished). **H.** Expression of DRA.R8-Gal4 exclusively in wild type DRA.R8 cells (arrow). H’ shows single GFP channel. **I.** Expression of DRA.R8-Gal4 spreads to most R8 cells in GMR-Hth flies. I’ shows single GFP channel. **J.** Distribution of R8 presynaptic sites visualized with UAS-brp^D3^:mKate, in wild type non-DRA R8 cells (left), wild-type DRA.R8 cells (middle), and R8 cells outside the DRA proper in GMR-Hth flies (right). **K.** Double labeling of R7+R8 presynaptic sites (rh3-Gal4 > UAS-brp^D3^:GFP) and DRA.R8 presynaptic sites (DRA.R8-LexA > LexAop-brp^D3^:mKate2), allowing for indirect extraction of DRA.R7 presynaptic sites (right, green). **L.** Summary of presynaptic site distribution between cell types. EM data for R7 (dark green) and R8 (dark red) was normalized to fit with brp^D3^ data. Note the difference between wild type DRA.R8 presynaptic site distribution (violet) and R8 presynaptic site distribution in GMR-Hth flies (black). Scale bar: 7μm in (D) for (D-E)

Polarization-sensitive neurons have been described in the central brain of several species, resulting in a powerful network model for how e-vector orientations represented in the central brain of insects ^34–36^. In contrast, very little is known about the anatomy of cell types directly post-synaptic to polarization-sensitive photoreceptors and the network architecture of medulla columns located in the DRA region ^37, 38^. For instance, it is unknown whether DRA columns accommodate the same or a different number of cellular units in order to perform their specific function (comparing e-vectors instead of wavelengths). Furthermore, it is unknown whether a given columnar cell type manifests DRA-specific morphological specializations, including differences in synaptic distribution and connectivity. As a minimal hypothesis, one could assume that DRA columns can very well serve skylight navigation by employing an invariant protocircuit that is identical to those columns processing color. Yet if modality-specific inter-column differences were to exist, they should be apparent at the DRA/non-DRA boundary, especially for cell types contacting photoreceptors from several neighboring ommatidia, like the distal medulla cell type Dm8, an amacrine-like cell type located in layer M6 where each cell collects information from a field of ∼14 R7 cells ^22, 39, 40^. We, therefore, asked whether Dm8 cells mix color and polarized light information by pooling photoreceptor inputs across the DRA boundary, when located at the dorsal edge of the medulla. Based on the very few examples of cellular differences between columns receiving input from pale and yellow ommatidia ^21, 22^, it seemed plausible to assume that specific differences could also exist between DRA and non-DRA medulla circuitry. We will refer to such differences as ‘modality-specific’ anatomical variations, since linearly polarized light, and not color, is processed by R7 and R8 and their downstream targets in this part of the visual system. This definition, therefore, centers on the physical nature of the stimulus and not on what the animal perceives as a modality ^41^.

In this study, we provide the first detailed neuroanatomical characterization of neuronal elements that are post-synaptic to R7 and R8 photoreceptor cells in the DRA region of the medulla of *Drosophila*. Using genetic redesign of the retinal mosaic, activity-dependent ‘GFP Reconstitution Across Synaptic Partners (GRASP) ^12^ and the trans-synaptic tracer ‘trans-Tango’ ^13^ we show that DRA.R7 and DRA.R8 cells are synaptically connected to distinct, newly-discovered Dm-DRA subtypes (Dm-DRA1 versus Dm-DRA2, respectively). These cells stratify in similar sublayers of M6 and specifically contact polarization-sensitive inputs, while avoiding contacts with color-sensitive non-DRA inputs. To achieve this DRA-specific connectivity, polarization-sensitive DRA.R8 cells differ from their color-sensitive counterparts both in layer targeting and the distribution of their presynaptic sites. Finally, using live imaging of ingrowing photoreceptor axons ^42^, we visualize the temporal sequence of R7 and R8 axons targeting to the M6 layer, thereby temporally separating synaptic partner choices. Our work reveals an instance for how repetitive microcircuits, organized in highly stereotypical medulla columns, can become reorganized locally in order to adapt them to their modality-specific needs. This newly described DRA circuit including a duplication of R7→Dm connections represents a perfect adaptation for comparing orthogonal angles of monochromatic, linearly polarized skylight with an equal synaptic weight.

## Results

### Presynaptic properties of DRA inner photoreceptors R7 and R8

In a first step towards characterizing DRA-specific differences in neural circuitry, we visualized inner photoreceptor axon terminals using the stochastic labeling technique ‘MultiColor FlpOut’ (MCFO) ^11^. We found that the typical layer-specific targeting of R7 (to layer M6) and R8 (to M3) observed across the medulla ^19^ is different in DRA columns, where both R7 and R8 terminate in M6, with one cell always terminating slightly more distally (Figure 1D). In agreement with previous reports ^32^, we found that this layer targeting pattern could be ectopically induced by transforming all ommatidia into the DRA fate in flies via over-expression of the transcription factor Homothorax in all photoreceptors (GMR-Hth) ^26, 43^, (Figure 1E). Quantification of the distance between R7 and R8 terminals in the medulla revealed a sharp boundary between DRA-specific and non-DRA layer targeting of inner photoreceptor cells (Figure 1F). In these experiments, R7 and R8 cell fates could not be distinguished, although we suspected DRA.R8 terminating distally from DRA.R7. We therefore generated a new driver expressed exclusively in DRA.R8 cells (DRA.R8-Gal4), using a mutated version of the *rh3* promoter in which three copies of a palindromic Prospero-binding repressor site from *rh5* and *rh6* promoters ^44^ had been inserted via site-directed mutagenesis (gift from Dmitri Papatsenko, unpublished; see materials & methods) (Figure 1G). Indeed, expression of this DRA.R8 reporter was absent in all R7 cells, resulting in DRA.R8-specific expression (Figure 1H), always labeling the DRA inner photoreceptor cell that terminated in a slightly more distal sublayer (Figure 1H; inset). In a GMR-Hth background, expression of DRA.R8-Gal4 expanded into most (if not all) R8 cells, albeit at varying expression levels (Figure 1I). We suspected that DRA-specific differences of R8 target layer selection could be indicative of modality-specific differences in post-synaptic circuitry. We first investigated the distribution of pre-synaptic sites in DRA-photoreceptors using transgenic fusions of the D3 domain of the active zone protein Bruchpilot (Brp) with fluorescent proteins ^45 46^. Indeed, its localization differed significantly between DRA.R8 cells and non-DRA R8 cells (Figure 1J). In contrast, the distribution of presynaptic sites from DRA.R7 cells (extracted indirectly via two-color labeling; see materials & methods), revealed a great resemblance between DRA.R7 and non-DRA R7 cells (Figure 1K). Interestingly, DRA.R8 cells manifested a more R7-like distribution of presynaptic sites, i.e. a significant shift of presynaptic sites towards the axonal tip (Figure 1L). Non-DRA R8 cells in GMR-Hth flies behaved quite differently (Figure 1J,L), with two distinct peaks clustered in M3 and M6. Importantly, the relative density distributions of fluorescent signals from non-DRA R7 and R8 presynaptic sites were in good agreement with EM data previously published ^23^ when normalized for comparison (see materials & methods).

### Modality-specific distal medulla cells at the dorsal edge of the medulla

To characterize the post-synaptic partners of R7 and R8 cells in layer M6 of the DRA region of the medulla, we focused on Dm8 cells, previously shown to be the main R7 target in non-DRA columns ^11, 22, 23, 39, 40^. Using the Dm8-specific driver GMR24F06-Gal4 and MCFO ^11^ we classified membrane contacts as potential sites for synaptic connections, whereas absence of contacts indicated absence of synaptic connections (see Supplemental Movie 1 and Materials & Methods). Morphology of Dm8-like cells at the dorsal pole of the medulla (Figure 2A, red and green) differed from adjacent Dm8 cells (blue) in that photoreceptor contacts were restricted to DRA inner photoreceptor terminals, while non-DRA photoreceptors were specifically avoided (Figure 2B). Furthermore, polar Dm8-like cells formed characteristic processes in a layer below M6 (termed ‘deep projections’, DP). Since these morphological features are never observed for non-DRA Dm8 cells ^11, 18^, we named this new cell type Dm-DRA1. Presynaptic sites were also detected on DPs, suggesting a potential role in communication between Dm-DRA1 cells and non-DRA cell types (Figure 2C). Interestingly, Dm-DRA1 cells manifested a considerable morphological diversity across the DRA (Figure 2D): Equatorial Dm-DRA1 cells were slimmer and more elongated and had less (posterior) to few short DP’s (anterior) compared to polar Dm-DRA1 cells. Irrespective of position, photoreceptor contacts of all Dm-DRA1 cells analyzed were restricted to DRA terminals (Figure 2 D’, red balls), hence making them modality-specific. Photoreceptor contact number was independent of a cell’s location (Figure 2E; Supplemental Figure S2A). Importantly, length and width of Dm-DRA1 cell clones within layer M6, as well as the length of the most prominent DP changed gradually as a function of the cells’ location along the DRA (Figure 2F-H)(Supplemental Figure S2B,C), rather than forming two distinct cell populations (equatorial vs polar). Importantly, clones resembling Dm-DRA1 cells were never observed at the ventral rim of the medulla (Figure 2 I). Previously, non-DRA Dm8 cells were shown to heavily overlap, therefore sharing many photoreceptor inputs ^22,39 23^. We found that non-DRA Dm8 cells never crossed the DRA boundary therefore never sharing photoreceptor contacts with Dm-DRA1 cells (Figure 2K). The average number of photoreceptor contacts were indistinguishable for non-DRA Dm8 and Dm-DRA1 clones (Figure 2L), a notable exception being Dm-DRA1 cells located close to the equator with a below-average number of contacts (Figure 2M). We, therefore, concluded that a strict modality-specific separation between color-specific Dm8 cells and polarization-specific Dm-DRA1 cells exists within layer M6 of the medulla.

**Figure 2:**
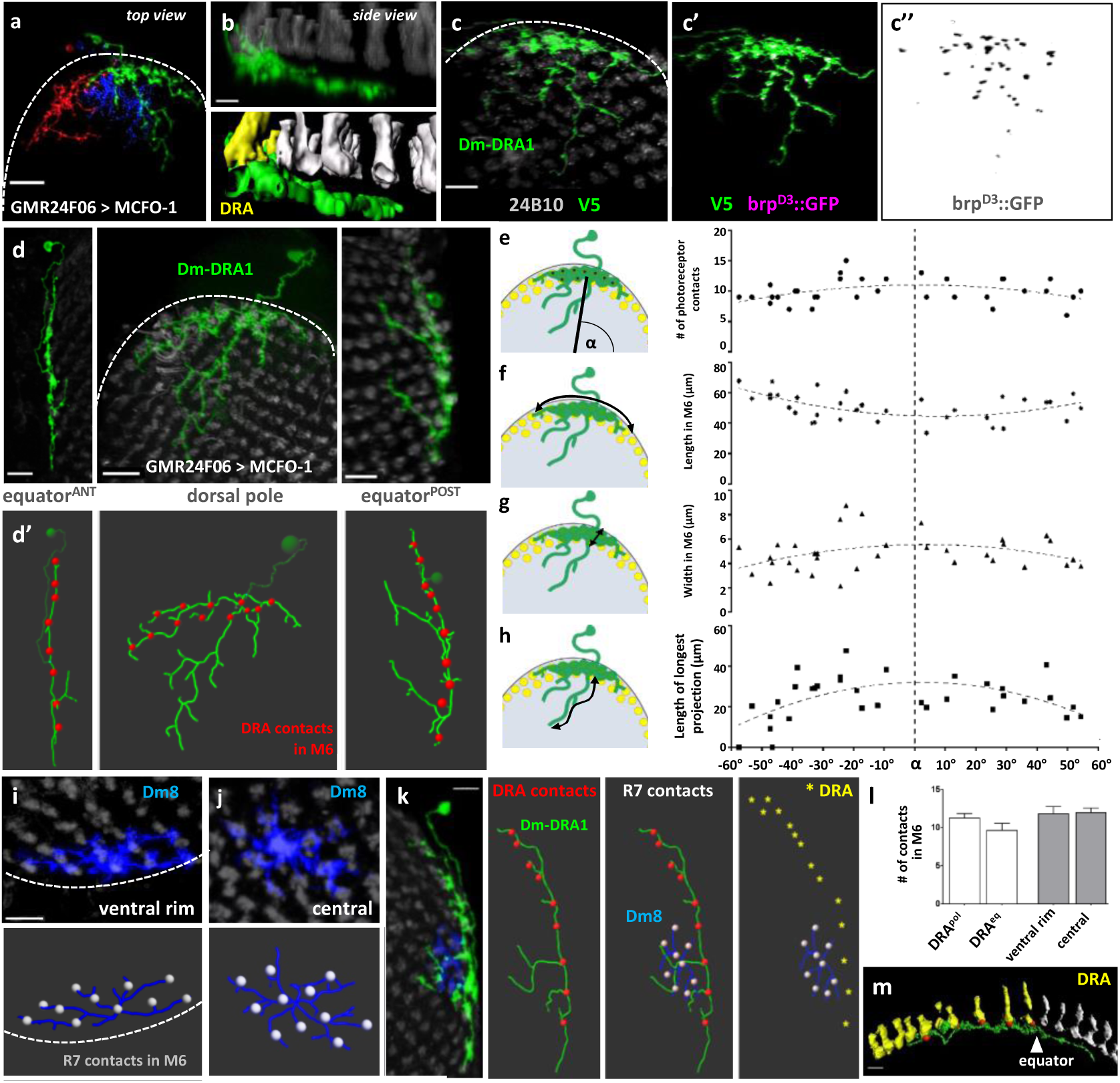
Morphology of modality-specific Dm-DRA1 cells at the dorsal edge of the medulla. **A.** Adult whole-mounted brain with MCFO ^11^ using Dm8-specific driver GMR24F06-Gal4 containing two clones touching the dorsal edge of the medulla (red cell and green cell). Dashed line is showing the edge of medulla. **B.** Top: Side view of green cell from (A). Note photoreceptor contacts are restricted to DRA photoreceptors (yellow in surface view, bottom), whereas contacts with non-DRA photoreceptors (grey) are avoided, resulting in ‘deep projections’ of the Dm-DRA1 cell penetrating the medulla centripetally, below the photoreceptors. **C.** Single cell clone of a polar Dm-DRA1 cell (green) co-labeled with UAS-brp^D3^::GFP ^45^ (purple), revealing presynaptic sites across the cellular surface, including the ‘deep projections’. **D.** Top: Three representative MCFO clones of Dm-DRA1 cells at different locations along the DRA: at the anterior equator (left), at the dorsal pole (middle), and at the posterior equator (right). Bottom (D’): Skeletons reconstructed from the above cells (photoreceptor contacts are visualized as red balls). Less ‘deep projections’ are present at the equator, especially anteriorly. **E.** Quantification of photoreceptor contact number per Dm-DRA1 cell (right) as a function of their position along the DRA (defined as Angle α drawn from the center of the medulla to the center of the Dm-DRA1 cell (left). **F, G.** Quantification of the length (F) and width (G) of a given Dm-DRA1 cell within layer M6 as a function of their position along the DRA. **H.** Length of the most prominent ‘deep projection’ for Dm-DRA1 clones as a function of their position along the DRA. **I.** Top: MCFO clone of a non-DRA Dm8 cell at the ventral rim of the medulla (top, blue). Bottom: skeleton and photoreceptor contacts as silver balls. Dashed line is showing the edge of medulla **J.** Top: MCFO clone of a central, non-DRA Dm8 cell clone (blue) with skeleton; photoreceptor contacts as silver dots (bottom). **K.** Representative whole mounted brain with adjacent MCFO clones of a Dm-DRA1 cell (green) and a non-DRA Dm8 cell (blue). These two cell types never share photoreceptor contacts, as visible from their skeletons (Dm8 contacts: silver balls; Dm-DRA1 contacts: red balls; yellow asterisks: DRA columns). **L.** Quantification of M6 photoreceptor contacts of Dm-DRA1 (polar and equatorial), and non-DRA Dm8 cells. **M.** Side view of a Dm-DRA1 surface rendering (green) at the equator. Note contacts (red balls) are restricted to DRA photoreceptor terminals (yellow), resulting in only 5 contacts for this cell. Scale bars: 15μm in (A); 5μm in (B), 7μm in (C),(D),(I),(J); 10μm in (K and M).

### Genetic re-design of the retina specifically alters Dm-DRA1 morphology

Due to their characteristic morphology, we suspected that Dm-DRA1 cells might represent a specialized cell type. In order to test this hypothesis, we repeated the MCFO-based morphological description using GMR24F06-Gal4 in a GMR-Hth background resulting in rather dramatic morphological changes for Dm-DRA1 cells (Figure 3A). Most notable was the strong increase in photoreceptor contacts: On average, a given Dm-DRA1 cell now contacted 25 incoming inner photoreceptor cells (as opposed to an average of 10, in wild type flies, Figure 3D). Our morphometric characterization revealed that the length of Dm-DRA1 cells in layer M6 remained unchanged when compared to wild type, but they now extended further into the medulla (width in M6; Figure 3B). Interestingly, morphology and photoreceptor contact number of non-DRA Dm8 cell clones remained indistinguishable from wild type (Figure 3C,D; Supplemental Figure 3B,C). Since Dm-DRA1 cells were affected in a GMR-Hth background, we asked whether Dm-DRA1 and non-DRA Dm8 cells now shared photoreceptor contacts, a situation never observed in wild type brains. Indeed, Dm8 cells and Dm-DRA1 cells now shared photoreceptor contacts due to the extended morphology of Dm-DRA1, ranging from 1 shared photoreceptor contact (posterior) to eight (anterior) (Figure 3E). Interestingly, non-DRA Dm8 cell clones appeared morphologically unaffected in GMR-Hth flies. However, quantification of GFP-positive Dm8 cell body numbers (including Dm-DRA cells) both in wild type flies and in a GMR-Hth background, revealed that about 50% of Dm8 cells were lost in the latter genotype (Figure 3F). Very similar results were obtained with another Dm8-specific driver, ort^C2b^-Gal4 (Supplemental Figure 3E, 3F).

**Figure 3:**
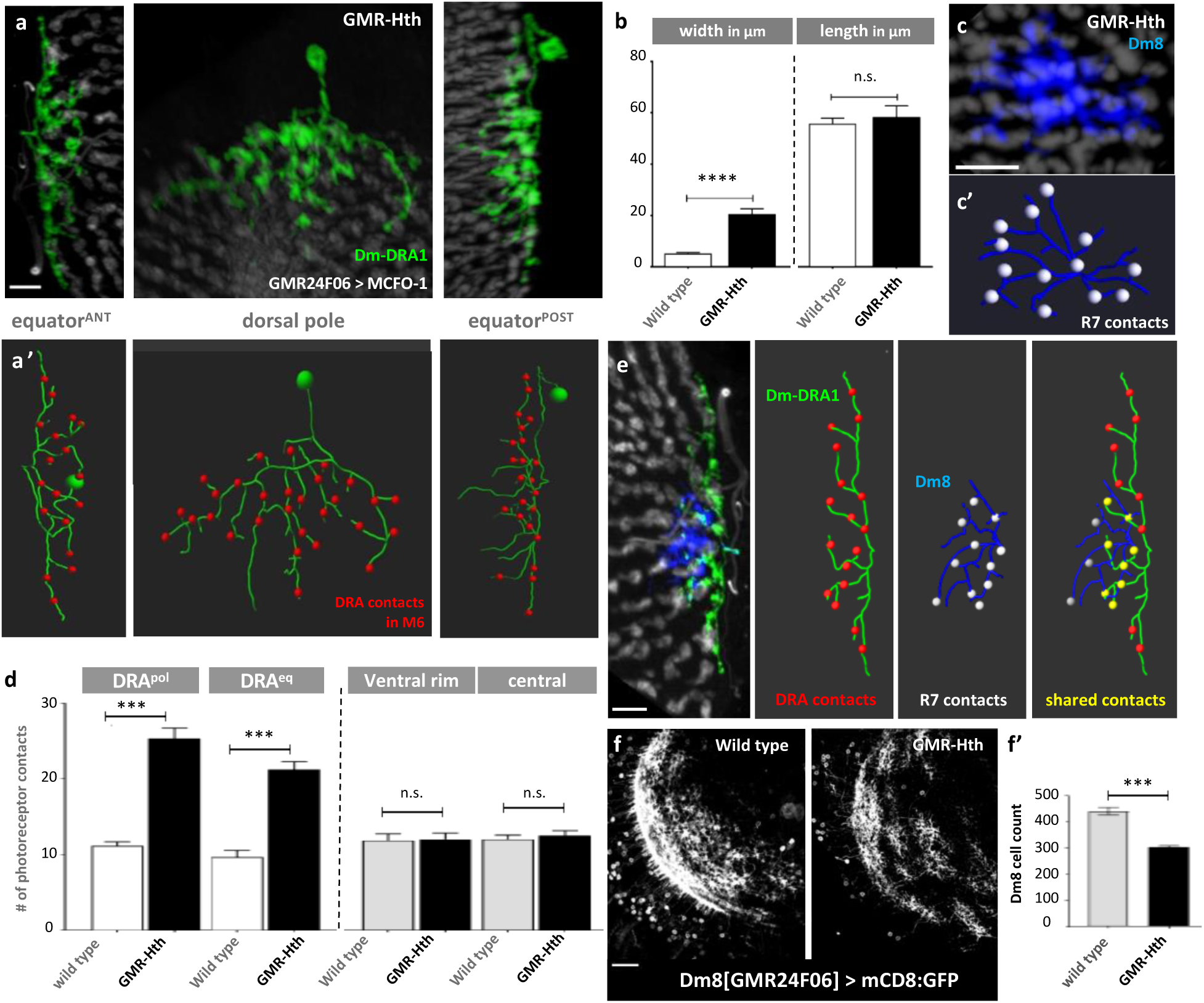
Genetic re-design of the retina specifically alters Dm-DRA1 morphology. **A.** Top: Three representative MCFO ^11^ clones of Dm-DRA1 cells labeled with Dm8-driver GMR24F06-Gal4 at different locations along the DRA in a GMR-Hth background: at the anterior equator (left), at the dorsal pole (middle), and at the posterior equator (right). Bottom (A’): Skeletons reconstructed from the above cells. Note the increase in photoreceptor contacts (red balls) and loss of ‘deep projections’. **B.** Comparison of length and width of Dm-DRA1 cells within layer M6 between wild type and GMR-Hth backgrounds. **C.** Representative MCFO clone of a non-DRA Dm8 cell (blue) in a GMR-Hth background, with skeleton (C’) and photoreceptor contacts (silver balls). **D.** Quantification of photoreceptor contact number across Dm cells, comparing non-DRA Dm8 cells and DRA-Dm1 cells in both wild type and GMR-Hth backgrounds. Note the increase of photoreceptor cell contacts in GMR-Hth is specific to Dm-DRA1 cells. **E.** Representative whole-mount brain with adjacent MCFO clones of a Dm-DRA1 cell (green) and a non-DRA Dm8 cell (blue) in a GMR-Hth background. The two cell types now share multiple photoreceptor contacts (yellow balls), as visible from their skeletons. **F.** Left: Single-channel images of whole mounted GMR24F06 > mCD8::GFP retinas from wild type flies (left) and GMR-Hth flies (right), depicting the reduced number of Dm8 cells. Right: Quantification of Dm8 cell number in wild type and GMR-Hth flies. Scale bars: 7μm in (A),(C),(E); 20μm in (F).

### Two types of modality-specific interneurons: Dm-DRA1 and Dm-DRA2

We confirmed the existence of modality-specific Dm-DRA1 cells using another, previously published Dm8 driver line (ort^C2b^-Gal4) ^22^, revealing the same gradual change in morphology from equatorial regions towards the dorsal pole (Supplemental Figure 4A). To our surprise, this driver line also labeled a second type of Dm cells that specifically contacted DRA inner photoreceptors but manifested morphological features significantly different from the Dm-DRA1 cells. These cells always lacked DPs (Figure 4A,B) and instead manifested ‘vertical projections’ (VP), processes that extend upwards, along the incoming inner photoreceptor terminals (Figure 4A’, 4B’), sometimes making photoreceptor contacts exclusively along the VP’s (Figure 4A’’, arrowheads; see Supplemental Movie 2). We, therefore, named this new cell type Dm-DRA2. Importantly, Dm-DRA2 always were modality-specific, i.e. they never overlapped or shared photoreceptor contacts with non-DRA Dm8 cells (Figure 4C). Furthermore, quantification of morphometric characteristics as a function of position along the DRA revealed no statistically significant variation of photoreceptor contacts (Figure 4D). Width and length of Dm-DRA2 cells gradually changed as a function of their position along the DRA, from slim and elongated at the equator, to more compact at the dorsal pole (Figure 4E,4F).

**Figure 4:**
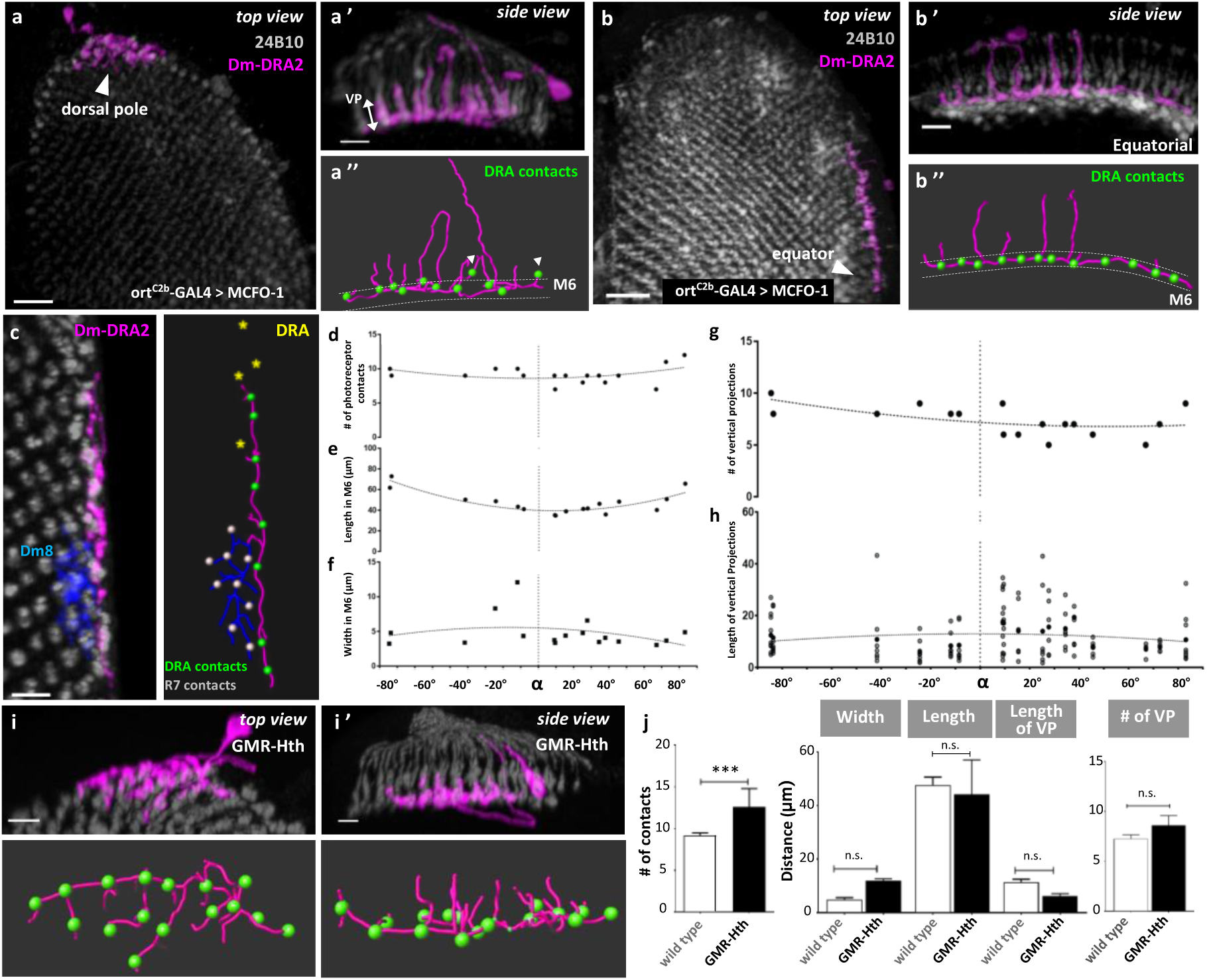
A second subtype of Dm-DRA cells with a different morphology. **A.** Adult whole-mounted brain with MCFO ^11^ experiment using Dm8-specific driver ort^C2b^-Gal4 revealing a morphologically different DRA-specific Dm cell type (termed Dm-DRA2) touching the dorsal edge of the medulla (purple). Note the absence of ‘deep projections’. Side view in A’ reveals unusual vertical projections along photoreceptor shafts. A’’: Corresponding skeleton with photoreceptor contacts (green balls). **B.** Representative MCFO clone using ort^C2b^-Gal4 containing a similar Dm-DRA2 cell (purple) with vertical projections (B’,B’’) located near the anterior equator. **C.** Two adjacent MCFO clones of a Dm-DRA2 cell (purple) and a non-DRA Dm8 cell (blue). These two cell types never share photoreceptor contacts, as visible from their skeletons (Dm8 contacts: white balls; Dm-DRA2 contacts: green balls; yellow asterisks: adjacent DRA columns). **D.** Quantification of photoreceptor contact number per Dm-DRA2 cell, as a function of their position along the DRA. **E.** Quantification of the length of a given Dm-DRA2 cell within layer M6 as a function of their position along the DRA. **F.** Quantification of the width of a given Dm-DRA2 cell within layer M6 as a function of their position along the DRA. **G.** Total number of vertical processes of Dm-DRA2 cells as a function of their position along the DRA. **H.** Mean length (solid dots) and spread of length of vertical projections (open circles) of Dm-DRA2 cells as a function of their position along the DRA**. I.** Representative clone of a Dm-DRA2 cell at the dorsal pole of the medulla from a GMR-Hth fly. Skeleton of Top view reveals no dramatic increase of photoreceptor contacts (green balls). Side view in I’ reveals vertical projections. **J.** Comparison of morphometric characteristics measured for Dm-DRA2 cells from wild type and GMR-Hth genotypes. Scale bars: 20 μm in (A and B); 10 μm in (A’ and B’); 7μm in (C, I and I’).

Interestingly, quantification of VP number (Figure 4G) and VP length (Figure 4H, average and individual) per Dm-DRA2 cell did not correlate with the cell’s position along the DRA. Finally, the change of Dm-DRA2 morphology was rather mild in a GMR-Hth background (Figure 4G): The increase in photoreceptor contacts was rather weak but significant (increase from 9,13 ± 0,350 to 12,6 ± 0,98) (Figure 4J), resulting in a weak increase of the cells’ width in M6. Finally, the number of VP’s increased mildly, while their average length decreased (Figure 4J). Additionally, a third Dm8-specific driver (Dm8 [ortc^2b^∩ort^c1-3^]-LexA) ^22^ also labeled both Dm-DRA1 and 2 cell types (data not shown). We, therefore, concluded that both cell types described here were modality-specific, apparently coexisting in the DRA region of the medulla.

### Distribution of Dm-DRA1 and Dm-DRA2 across the DRA region of the medulla

Since Dm8 cells throughout the non-DRA region of the medulla heavily overlap and share photoreceptor contacts, we proceeded to investigate overlap of Dm-DRA1 and Dm-DRA2 cells within the DRA region. As expected, stochastic Dm-DRA2 clones heavily overlapped, sharing between 2 and 9 photoreceptor contacts (Figure 5A; Supplemental Figure 5E). An artificial overlay of skeletons from all characterized Dm-DRA2 cells revealed a dense coverage of the DRA region without DP’s but with numerous VP’s (Figure 5B, Supplemental fig 5B). A similar analysis for Dm-DRA1 also revealed heavy overlaps between their own kind (Figure 2K, Supplemental Figure 5C), with number of overlaps ranging from 2-7 (Supplemental Figure 5E). Importantly, the artificial overlay of all analyzed Dm-DRA1 skeletons revealed a significantly different arrangement of processes, with characteristic DP’s spreading centripetally from the dorsal pole of the medulla (Figure 5D, Supplemental Figure 5A). Since both Dm-DRA cell types cover the entire DRA region of the medulla, we suspected they co-exist at any given location within the DRA. Using ort^C2b^-Gal4 and MCFO, we, therefore, identified brains with clones of Dm-DRA1 and Dm-DRA2 cells in close apposition (Figure 5E). From a top view, it appeared that processes from both cell types were closely intermingled. However, when viewed from the side, it became apparent that the two Dm-DRA cell types tend to stratify in two slightly different sublayers within M6 (Figure 5E’). Importantly, the peak of Dm-DRA2 signal was always located more distally (Figure 5E’ and 5F). Given that only DRA inner photoreceptors manifest modality-specific layer targeting into different sublayers of M6 (R8 always terminating distally from R7 - for direct comparison of distances, see Figure 5G), we hypothesized that Dm-DRA1 and Dm-DRA2 could be specific targets of DRA.R7 and DRA.R8, respectively.

**Figure 5:**
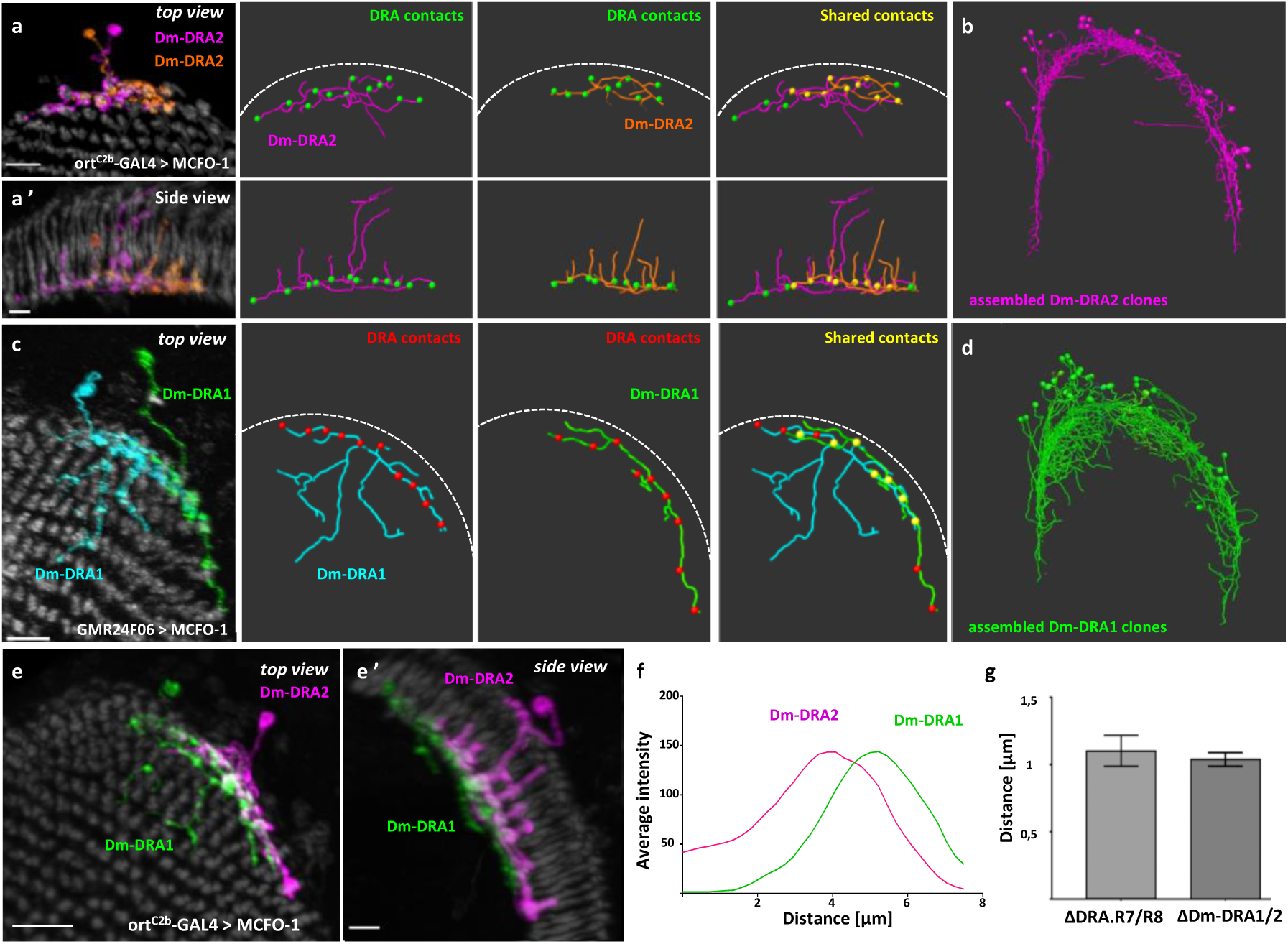
Both Dm-DRA cell types densely cover the DRA region of the medulla. **A.** Top: Representative whole mounted brain with two adjacent MCFO ^11^ clones of Dm-DRA2 cells (purple and orange, respectively). Skeletons depict photoreceptor contacts (green balls). The two cells extensively share DRA photoreceptor contacts (yellow balls). Bottom (A’): Side view of the same clones in A. **B.** Artificially generated assembly of all Dm-DRA2 MCFO clones analyzed, aligned onto a standard brain. Note the absence of ‘deep projections’ protruding centripetally. **C.** Two adjacent MCFO clones of two Dm-DRA1 cells (green and cyan, respectively). Skeletons depict photoreceptor contacts (red balls). The two cells extensively share contacts (yellow balls). **D.** Assembly of all Dm-DRA1 clones analyzed (generated as in B). **E.** Two adjacent Dm-DRA cell clones of different subtypes (Dm-DRA1: green; Dm-DRA2: purple) located at the same position along the DRA. E’: side view. **F.** Dm-DRA layering reveals positioning of Dm-DRA1 cells in close proximity, yet the peak of their signal is always distally from Dm-DRA2 cells. **G.** Comparison of distance between R7 and R8 terminals in the DRA with distances measured between Dm-DRA1 cell layer and Dm-DRA2 cell layer. Scale bars: 7 μm in (A and A’); 10 μm in (C),(E and E’).

### Synaptic connectivity between inner photoreceptor subtypes and Dm-DRA cells

In order to test which photoreceptors are synaptically connected to either Dm-DRA subtype, we first used activity-dependent ‘GFP Reconstitution Across Synaptic Partners’ (GRASP) ^12, 47^. First, we tested Dm-DRA1 connectivity using GMR24F06-LexA, which (like its Gal4 counterpart) is never expressed in Dm-DRA2 cells. GRASP signals with all photoreceptors (longGMR-Gal4) were restricted to layer M6, across the medulla (Figure 6A). In contrast, no GRASP signal was obtained with DRA.R8-Gal4 (Figure 6B). These experiments were repeated with [ort^c2b^∩ ort^c1-3^]-LexA, expressed in both Dm-DRA1 and Dm-DRA2. Once again, GRASP signal with all photoreceptors extended across medulla layer M6 (Figure 6C). Interestingly, GRASP signal extended far more vertically in the DRA, along the incoming shafts of polarization-sensitive photoreceptors. Importantly, virtually identical, long DRA-specific vertical extension of the GRASP signal was obtained using DRA.R8-Gal4, suggesting these signals originate from DRA.R8 → Dm-DRA2 connections (Figure 6D). We, therefore, compared the total vertical length of DRA GRASP signals with the vertical depth of either Dm-DRA1 or Dm-DRA2 cell signals, as well as with the vertical spread of DRA.R7 versus DRA.R8 presynaptic sites (Figure 6E,F). The photoreceptor GRASP signals obtained with Dm-DRA1 matched the length of its vertical depth, as well as the most terminal peak of DRA.R7 presynaptic sites (Figure 6E). Conversely, the DRA GRASP signal between DRA.R8 and Dm-DRA1+2 cells matched the average length of Dm-DRA2 VP’s and overlapped with the distribution of DRA.R8 presynaptic sites (Figure 6F). We then expressed the trans-synaptic tracer ‘trans-Tango’ ^13^, using the very specific DRA.R8-Gal4 (Figure 6G). Sparse labeling (see materials & methods) revealed single cells with strikingly resembling Dm-DRA2 cells, as identified by the absence of DP’s and the presence of VP’s (Figure 6H). Under saturating conditions the entire DRA region of the medulla was labeled (Figure 6I). Importantly, this signal did not include any DP’s (particularly visible around the dorsal pole), and instead manifested multiple VPs, therefore resembling the artificial overlay of Dm-DRA2 skeletons from Figure 4B, while lacking the characteristic signals from the Dm-DRA1 overlay. Finally, forcing all R8 cells to layer M6 in GMR-Hth flies resulted in photoreceptor → Dm8 GRASP signals in M6 ^33^(Figure 6 J,K). We, therefore, concluded that only in the DRA, both R7 and R8 cells are synaptically connected to different, morphologically distinct Dm-DRA subtypes.

**Figure 6:**
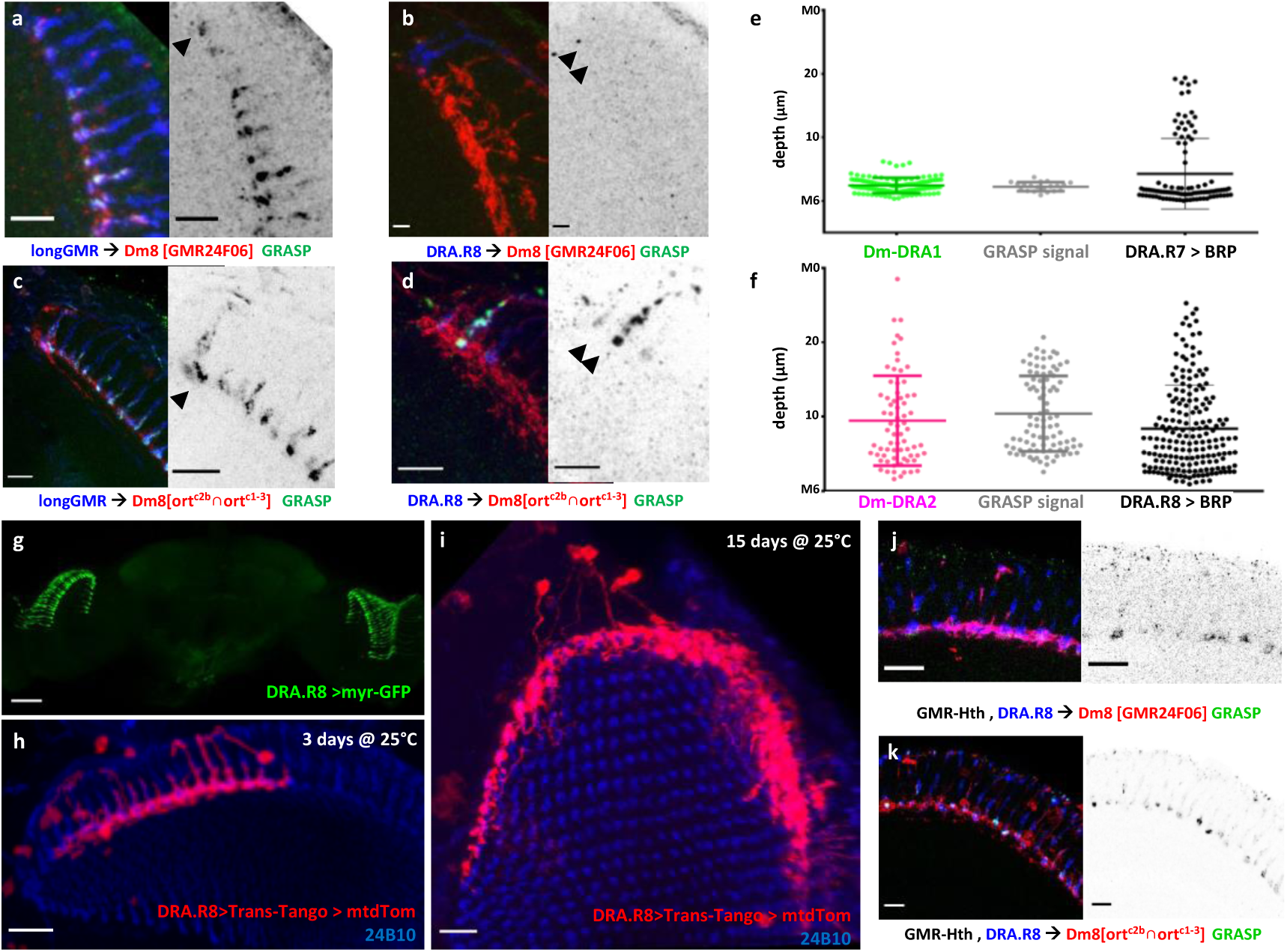
GRASP and trans-Tango reveal R7- and R8-specific Dm-DRA subtypes. **A.** Activity GRASP experiment visualizing potential synaptic contacts between photoreceptors (labeled with longGMR-Gal4, UAS-nsyb:spGFP^1–10^) and Dm-DRA1 cells (labeled with GMR24F06-LexA, LexAop-CD:spGFP^11^). GRASP signal is restricted mostly to layer M6 throughout the medulla, including the DRA. **B.** Absence of activity GRASP between DRA.R8 photoreceptors (labeled with DRA.R8-Gal4, UAS-nsyb:spGFP^1–10^) and Dm-DRA1 cells (labeled with GMR24F06-LexA, LexAop-CD:spGFP^11^). **C.** Activity GRASP between photoreceptors (labeled with longGMR-Gal4, UAS-nsyb:spGFP^1–10^) and DmDRA1+2 cells (labeled with ort^c2b^ ∩ ort^c1-3^-LexA, LexAop-spGFP^11^). GRASP signal spreads vertically, especially in the DRA. **D.** Activity GRASP between DRA.R8 photoreceptors (labeled with DRA.R8-Gal4, UAS-nsyb:spGFP^1–10^) and Dm-DRA1+2 cells (labeled with ort^c2b^ ∩ ort^c1-3^-LexA, LexAop-CD:spGFP^11^). Note vertical extension of the GRASP signal. **E.** Quantitative comparison of vertical extension of Dm-DRA1 cells, activity GRASP signal from (A) and distribution of DRA.R7 presynaptic sites (from Figure 1). **F.** Quantitative comparison of vertical extension of Dm-DRA2 cells, activity GRASP signal from (D) and distribution of R8 presynaptic sites (from Figure 1). **G.** Expression of DRA.R8-Gal4 from brain used for trans-Tango tracing ^13^. **H.** Sparse trans-Tango tracing (3 days, 25°C degrees) reveals cells resembling Dm-DRA2 (note presence of vertical projections and absence of deep projections). **I.** Dense trans-Tango experiment (15 days, 25°C degrees) reveals processes covering the entire DRA (note absence of deep projections, as well as presence of vertical projections). **J.** Activity GRASP ^12^ between DRA.R8 photoreceptors (labeled with DRA.R8-Gal4, UAS-nsyb:spGFP^1–10^) and Dm-DRA1 cells (labeled with GMR[24F06]-LexA, LexAop-CD4:spGFP ^11^) in GMR-hth flies. Note GRASP signal is restricted to M6. **K.** Activity GRASP between DRA.R8 photoreceptors (labeled with DRA.R8-Gal4, UAS-nsyb:spGFP^1–10^) and DmDRA1+2 cells (labeled with ort^c2b^ ∩ ort^c1-3^-LexA, LexAop-CD4:spGFP^11^) in GMR-hth flies (note GRASP signal is restricted to M6). Scale bars: 7 μm in (A-D) and (J and K); 50 μm in (G); 10 μm in (H and I).

### Sequential targeting of DRA.R7 and DRA.R8 photoreceptor terminals to layer M6

Layer targeting of R7 and are R8 cells to medulla layers M6 and M3, respectively, is a well-understood process ^48^. Furthermore, pre-sorting of synaptic partners in the appropriate layer may be critical to bring correct partners in spatial vicinity for synapse formation ^49^. For instance, forcing the termination of R8 cells in layer M6 will lead to the formation of synapses between R8 and Dm8 cells, which are never observed in wild type flies ^33^, and even redirecting R1-R6 in the medulla leads to synapses with four postsynaptic partners, typical for R1-R6, even though the correct postsynaptic neurons are not present ^50^. We, therefore wondered how DRA.R7 and DRA.R8 cells could reliably choose between different post-synaptic partners when terminating in such close proximity within their respective M6 sublayers. In order to get some insight into this process, we decided to characterize the time series of DRA.R8 photoreceptor axons growing into the medulla, using 2-photon live imaging ^42^(see material and methods) (Figure 7A, Supplemental Movie 3). Imaging sparsely labeled R7 and/or R8 cells (within the DRA and outside of it) from an *ex vivo* brain explant over the course of 18 hours revealed the temporal dynamics of their individual outgrowth behavior during layer targeting (Figure 7B). Most strikingly, around ∼43-47% pupation DRA.R8 cells originally behave like ‘normal R8 cells, i.e. pausing at the distal border of the medulla, sending thin filopodia exploring prospective layer M3 After 47% pupation, DRA.R8 cell growth cone in non-DRA R8 recipient layer began to thicken, while simultaneously growing towards the deeper R7 recipient layer. Such behavior was never observed in non-DRA R8 cells, which still elongated after reaching the non-DRA R8 recipient layer, most likely via a passive process as the medulla thickens ^42, 51, 52^. The speed and dynamics of the thickened DRA.R8 process descending towards M6 suggest it results from an active process. Taken together, these data show that despite their striking similarity, R7 and R8 cells in the DRA arrive sequentially at their destination in M6, thereby separating their ability to contact postsynaptic partners in time.

**Figure 7:**
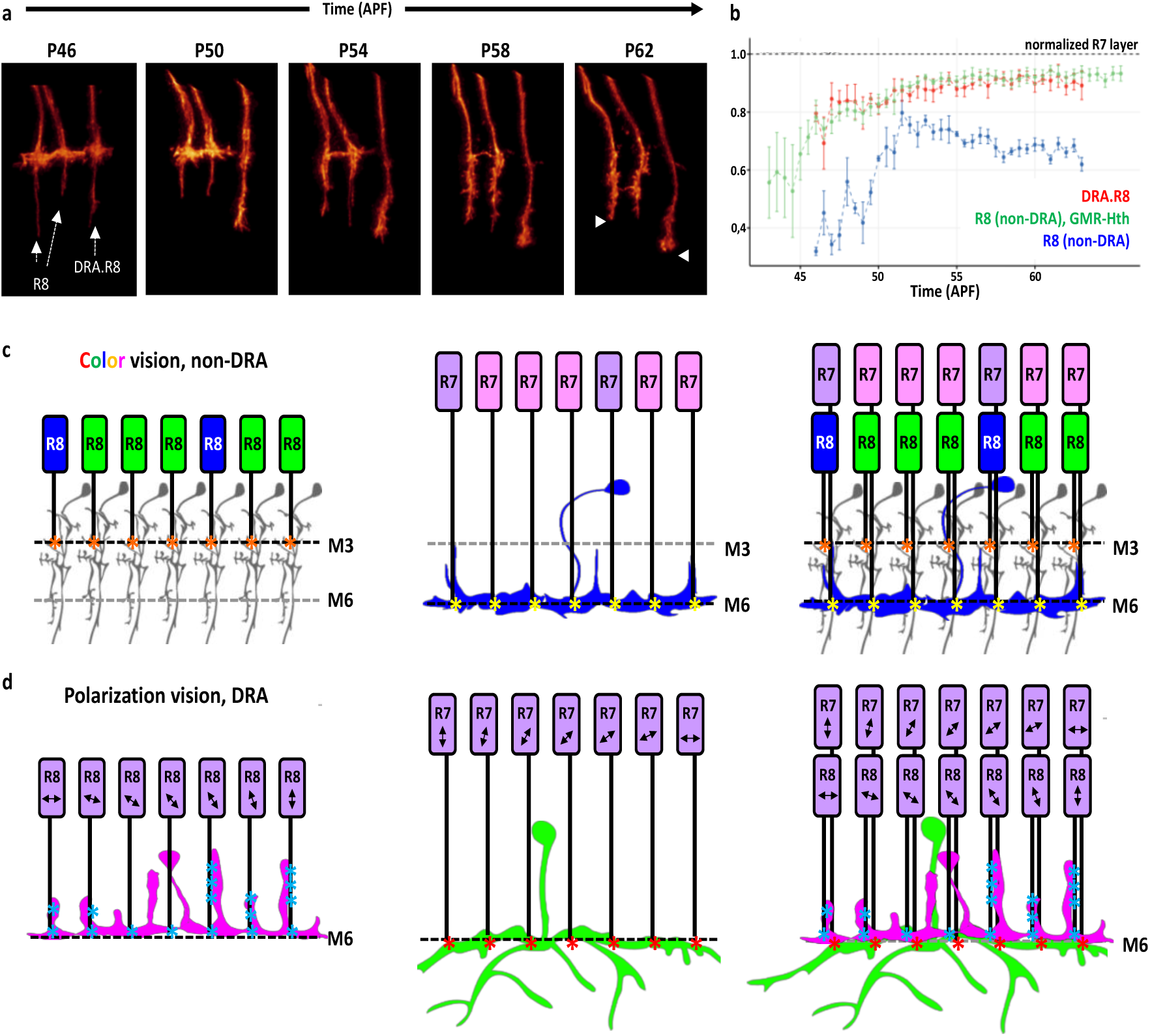
Temporal dynamics of DRA.R8 cells reaching layer M6. **A.** Live imaging of DRA.R8 layer targeting in *ex vivo* brain cultures ^42^. Note stabilization, thickening and extension of filopodium reaching towards M6. **B.** R8 temporal dynamics normalized with regard to the position of DRA.R7 (dashed line). Comparison of non-DRA R8 (blue), wild type DRA.R8 (red) and R8 cells in GMR-Hth flies (green). **C.** Summary model depicting color-sensitive R7 and R8 photoreceptor layer targeting and their main post-synaptic partners: Tm5 cells (grey) are columnar and post-synaptic to R8 cells in M3 (synapses symbolized by orange asterisks); Dm8 cells (blue) are multicolumnar and post-synaptic to R7 cells in M6 (synapses symbolized as yellow asterisks). **D.** Summary model depicting DRA circuitry: Dm-DRA2 cells (purple) are multicolumnar, form vertical projections and are post-synaptic to DRA.R8 cells (synapses symbolized by light blue asterisks); Dm-DRA1 cells (green) are also multicolumnar, form deep projections under color-sensitive photoreceptors and are specifically post-synaptic to DRA.R7 cells in M6 (synapses symbolized as red asterisks). Note that Dm-DRA2 cells are located slightly more distally within layer M6.

## Discussion

Here we describe specific differences in the wiring diagram of DRA medulla columns. There, both R7 and R8 provide UV inputs that differ only by their e-vector information. This represents a striking difference to the color vision system in non-DRA columns, where Dm8 cells collect from ∼14 R7 cells, whereas the main post-synaptic partner of R8 cells are columnar Tm5c cells (Figure 7C). The latter circuit provides the anatomical basis for strong preference of UV inputs over longer wavelengths, as observed behaviorally ^22, 40^. Our activity GRASP experiments also revealed connections between DRA photoreceptors and Tm5c cells, albeit weaker than in non-DRA columns (Supplemental Figure S6C), suggesting these connections still exist and polarized light information may be transmitted to the lobula neuropil via Tm5c-like cells. In agreement with this model, our trans-Tango experiments using DRA.R8-Gal4 also reveal columnar medulla cells types, whose cell fate could not be determined (Supplemental Figure S6 E,F). Most strikingly, we show that in the DRA, both R7 and R8 inputs (i.e. orthogonal e-vector orientations) are processed separately, yet using similar post-synaptic elements (Dm-DRA1 versus Dm-DRA2). This circuit design appears ideal for assigning an equal synaptic weight to both signals to be compared (similar e-vector analyzer directions from neighboring ommatidia) (Figure 7D).

### The DRA.R8 cell resembles an R7 cell in several ways

R8 cells in the DRA manifest several features normally only found in R7, like expression of Rh3 and layer-specific targeting to M6, instead of M3. We show that in wild type flies, DRA.R8 always terminates slightly distally from DRA.R7, with their terminal thereby forming two distinct sublayers within M6. Interestingly, layer M6 has previously been subdivided into M6a and M6b, based on slight differences in (non-DRA) R7 layer targeting ^18^. Although the sublayers we observed in the DRA are very similar to M6a and M6b, this could not be confirmed due to the lack of molecular markers distinguishing between them. Instead, we asked whether layer M3 within the DRA still manifests its most prominent, previously described anatomical hallmarks. Using previously published cell-type specific driver lines, we confirmed that laminar monopolar cell type L3 correctly terminates in M3 within the DRA, indistinguishable from non-DRA columns (Supplemental Figure 1B). Similarly, distal medulla cells types Dm4, Dm12, and Dm20 also stratify normally within M3 (Supplemental Figure S1C,D,E). It appears, therefore that layer M3 forms normally within the DRA, yet DRA.R8 cells efficiently bypass it, via a mechanism that remains unknown.

Over-expression of Hth is sufficient to induce most if not all cell-autonomous programs that lead to the morphological specialization of DRA.R8 cells ^26, 31^. In this genotype, all R8 cells are forced into layer M6, yet the very close and slight disorganization observed between R7 and R8 terminals could be due to constant over-expression of Hth inducing a higher variance in R8 layer targeting precision. Alternatively, it might be due to the fact that their correct post-synaptic target (Dm-DRA2) is missing in the main part of the medulla. In agreement with this model, we have never observed two morphologically distinct subtypes of Dm8-like cells outside the DRA. Deep projections, a hallmark feature of Dm-DRA1 cells, or long vertical projections (Dm-DRA2 cells), are never induced in Dm8 cells outside the DRA, neither in the wild type nor in flies over-expressing Hth in photoreceptors. Interestingly, the distribution of DRA.R8 presynaptic sites becomes bimodal in a GMR-Hth background (with peaks in M3 and M6), suggesting these cells might combine non-DRA synapses in M3 with new connections in M6. Our GRASP experiments in GMR-Hth flies show that transformed R8 cells now form synapses in layer M6 with those non-DRA Dm8 cells that have survived the transformation event (Figure 6J,K). Hence, it appears that in absence of Dm-DRA2 cells, genetically induced DRA.R8 cells combine synaptic connections that resemble those of wild type R7 and R8 cells.

### How to define a Dm8 cell - how many different subtypes are there?

Using different published Dm8-specific Gal4 and LexA driver lines, we characterized two different types of Dm-DRA cells with distinct morphologies. Interestingly, GMR24F06-Gal4 is expressed specifically in Dm-DRA1 cells (and never in Dm-DRA2), albeit at lower levels than in non-DRA Dm8 cells. This becomes particularly apparent upon co-labeling with the other Dm8 drivers (Supplemental Figure 4B). The two remaining Dm8 drivers (ort^C2b^-Gal4 and [ort^c2b^∩ ort^c1-3^]-LexA) are expressed in both Dm-DRA cell types at levels comparable to non-DRA Dm8 cells. The strong morphological differences between Dm-DRA cells and non-DRA Dm8 cells, between Dm-DRA1 and Dm-DRA2 types, or even within the Dm-DRA1 subtype (equatorial versus polar) raised the question whether all these cells are indeed Dm8 cells. The definition of an adult medulla cell type relies on stereotypical morphology across medulla cartridges. On one hand, there are similarities common to all these cells: Dm-DRA cells manifest the same number of photoreceptor contacts as non-DRA Dm8 cells irrespective of their location (only exception being Dm-DRA1 cells touching the equator). On the other hand, Dm-DRA cells show rather dramatic, modality-specific morphologies. Around the dorsal pole, Dm-DRA1 cells manifest prominent ‘deep projections’ (DP’s), i.e. processed that avoid contacts with non-DRA photoreceptors and stratify below the M6 layer. We show that these processes contain presynaptic sites, making it likely that Dm-DRA1 communicate with other, non-DRA specific cells whose identity remains unknown. These synaptic contacts could represent one of the first sites of integration between celestial compass information and chromatic information, or point light sources, like the sun ^53 54^. Dm-DRA2 cells have no DPs, instead, they form ‘vertical projections’ (VP’s) which form intimate contacts with photoreceptor shafts. Our GRASP experiments reveal that these VP’s are the likely site where synapses with DRA.R8 are made. Due to their unusual morphology, we systematically compared both Dm-DRA cells types with previously published cell types, to exclude the possibility that we misclassified them. For instance, Dm-DRA2 shows some resemblance with Dm11 cells, which also form long vertical projections ^11^. However, we excluded a misclassification, since MCFO experiments using Dm11-specific driver R11C05-GAL4 revealed the existence of Dm11 cells with the expected morphology within DRA columns (Supplemental Figure 7A,B,C,D). To be absolutely sure, we documented the DRA morphology for all Dm cell types for which drivers are available ^11^, and none of them resembled the Dm-DRA cell types we described here (data not shown). Our data therefore points towards two different Dm-DRA cell types co-existing exclusively in the DRA region of the medulla, where they replace Dm8 cells.

### Two Dm-DRA subtypes and the problem of wiring them

Both Dm-DRA cell types described here specifically contact polarization-sensitive inner photoreceptors, while avoiding photoreceptor contacts with non-DRA inputs. Both GRASP and trans-Tango together confirm our prediction that Dm-DRA1 cells are post-synaptic to DRA.R7, whereas Dm-DRA2 is post-synaptic to DRA.R8 (although we cannot rule out connections with DRA.R7 as well). The existence of two distinct Dm-DRA cell types being post-synaptic to DRA.R7 versus DRA.R8 make sense in the light of one single Dm8-like cell post-synaptic to both photoreceptors would be integrating signals with orthogonal e-vector tuning. Instead, Dm-DRA1 and Dm-DRA2 appear to process orthogonal signals, by collecting from neighboring ommatidia with slightly different e-vector analyzer directions. For now, it remains unknown whether any given Dm-DRA cell forms an equal number of synapses with each of the DRA.R7 (or DRA.R8) cells it contacts, or whether its inputs are dominated by one specific cell, for instance by concentrating the majority of synapses made within one column. In the former case, e-vector tuning of any Dm-DRA cells would be rather broad, due to an averaging over inputs from ∼10 neighboring ommatidia with gradually changing analyzer directions within the fan-shaped array of the DRA, whereas the latter case could result in sharper responses. The recently published whole brain dataset will hopefully reveal this crucial information ^55^. Unfortunately, nothing is known about the physiological responses of Dm8 cells, hence one can only speculate about Dm-DRA responses. One would assume such responses to be modulated sinusoidally, under a rotating pol filter, exhibiting phases of both hyper-and depolarization, as previously described for polarization-sensitive neurons in other insect species ^34^. It remains unclear how exactly these response properties arise, since both DRA.R7 and DRA.R8 are histaminergic and their outputs are already polarization-opponent in nature, as recently demonstrated ^29^. Although the ‘compass pathway’ leading to the representation of celestial e-vector orientations in the central complex via the anterior optic tubercle has been described in great detail in larger insects ^53^, comparably little data exists on polarization-sensitive medulla circuit elements and their circuitry ^38^. The two modality-specific Dm-DRA cell types described here seem well suited as the first-level elements of this pathway. One pressing question is at which level their orthogonal signals are compared. Their close proximity suggests there could be synaptic connections between Dm-DRA1 and 2, yet this cannot be tested due to the lack of specific drivers. Alternatively, their outputs could be processed separately by unknown post-synaptic targets.

DRA.R7 and DRA.R8 cells choosing separate post-synaptic partners seems like a difficult task, due to the close proximity of Dm-DRA1 and Dm-DRA2 in layer M6. Our live imaging data reveals that, despite their similarities in target layer selection, a temporal delay exists between ingrowing R7 versus R8 terminals, even in the DRA. The importance of tightly controlled temporal windows during which R7 and R8 establish specific connections has previously been demonstrated ^33, 56^. Over-expression of Hth clearly shows that both Dm-DRA cell types change morphology, while non-DRA Dm8 morphology is unaffected. This data points towards an unknown molecular signal present on DRA inner photoreceptors to be received by Dm-DRA cells to ensure modality-specific connectivity. It appears that in wild type flies, non-DRA Dm8 cells preferentially contact non-DRA photoreceptors, while avoiding contacts with DRA inputs. It was recently shown that immunoglobulin-family cell surface proteins of the DIP/Dpr families might play an important role in mediating such R7 → Dm8 contacts outside the DRA, in an ommatidial subtype-specific fashion ^57, 58^. It remains to be seen if similar proteins are at play in the DRA region of the medulla. Interestingly, non-DRA Dm8 cells in GMR-Hth flies will connect with both R7 and R8 now terminating in M6 (Figure 6J,K). However, we find that the total number of Dm8 cells is reduced, potentially due to Dm8 cells undergoing apoptosis either after encountering the ‘wrong’ photoreceptor terminals or after they fail to make sufficient number of synapses altogether.

### Two R7-like circuits in one column and their evolutionary implications

Due to the similarity between DRA.R8 cells and DRA.R7 cells (Rhodopsin expression, lack of Senseless, projections to M6, distribution of presynaptic sites, Dm8-like post-synaptic target), we propose that *Drosophila* DRA ommatidia contain two R7-like cells for processing information from orthogonal e-vector analyzers. This situation resembles the ommatidial design of some larger hymenopteran and lepidopteran insects, where every ommatidium contains two ‘true’ R7 photoreceptor cells which are specified early on during development from an uncommitted pool of precursor cells ^59^. In both species, the extra R7 cell is believed to enable a superior form of color vision, via stochastic choice between pale and yellow fates, resulting in three ommatidial subtypes, rather than two. It was recently shown that the butterfly homolog of Spineless is also necessary for pale/yellow choices in *Papilio*, suggesting the molecular mechanism is conserved between these distantly related species ^60^. Unlike these larger insects, fly ommatidia always contain only eight photoreceptor neurons and it is believed that a ninth one cannot easily be generated in the larval eye disc. We propose that flies have adopted a different strategy for obtaining a second, R7-like cell in DRA ommatidia, by transforming the R8 photoreceptor (i.e. the ommatidial founder cell) into an R7-like cell, beginning at mid-pupation: Expression of Senseless, a transcription factor crucial for specifying R8 cells ^61, 62^ is turned off ^26, 31^, and expression of an R7 Rhodopsin (Rh3) is induced soon after. Interestingly, our live imaging data reveals that DRA.R8 cells start descending towards layer M6 at exactly the same time (∼47% pupation), when Senseless is turned off ^26, 31^. Furthermore, we show that the distribution of presynaptic sites in a DRA.R8 cell also becomes significantly more R7-like, with a stronger clustering towards the tip of the photoreceptor cell. Furthermore, this second R7-like cell is equipped with its own Dm8-like (i.e. R7-specific) post-synaptic circuit element (Dm-DRA2). Our results can therefore serve as a model to address the larger questions of how signals from two ‘true’ R7 cells might be processed in bees and butterflies. Are the signals from R7 cells of the same ommatidium pooled in the same post-synaptic cell, or processed by two separate Dm8-like cells that pool over several ommatidia, like in the fly DRA? Sparse labeling of medulla cell types in bees have described several amacrine-like cell types, yet no clear Dm8-like candidates ^63, 64^. It therefore remains unknown whether two different types of Dm8 exist throughout the main part of the medulla, in these species. However, it should be noted that the two R7-like photoreceptors within honeybee ommatidia terminate in slightly different medulla sublayers ^65^, bearing striking resemblance to what we describe for the fly DRA. It is therefore possible that this segregation into different medulla sublayers also reflects differences in their post-synaptic Dm8-like circuitry. Furthermore, specific wiring differences between stochastically-distributed, color-sensitive columnar units in butterflies have been described, both for inter-photoreceptor connections ^66^, as well as axon collaterals within the lamina neuropil ^67^.

Taken together, our data reveal how the integration of morphologically and synaptically distinct cellular units into a specific subgroup of repetitive columnar microcircuits serves as the anatomical basis shaping their modality-specific function.

## Materials and methods

### Fly stocks and Fly rearing

The flies were maintained on standard molasses-corn food at 25°C 12h light/dark cycle incubator unless otherwise mentioned. The following flies were used in the study:

- *Drivers*: GMR-Gal4, GMR24F06 ^11^, ort^c2b^-Gal4 ^22^ and ort^C1-3^-LexADBD, ort^C2B^-dVP16AD ^22^; ort^C1a^-Gal4DBD#3, ok371-dVP16AD ^22^, GMR23G11-GAL4 (Bloomington stock center), GMR47G08-GAL4 (Bloomington stock center), GMR11C05-GAL4 (Bloomington stock center), *rh3*-Gal4^-137^ (gift from T. Cook), DRA.R8-Gal4 (this study), DRA.R8-LexA (this study), GMR-FRT-stop-FRT-Gal4 (Bloomington stock center), and sens-Gal4 (gift from R. Hiesinger).
- *MCFO*: MCFO-1^11^, tub***-***FRT***-***Gal80***-***FRT (gift from N. Özel), and hsFLP (Bloomington stock Center)
- *Active zone localization*: UAS-brp^D3^::GFP, UAS-brp^D3^::mKate2 and LexAop-brp^D3^::mKate (gift from Stefan Sigrist). *GRASP experiments*: UAS-nSyb-spGFP^1–10^, lexAop-CD4-spGFP^11,^ lexAop-nSyb-spGFP^1–10^ and UAS-CD4-spGFP^11,12^
- *Cell labeling*: UAS-IVS-myr::smGdP-V5, UAS-mCD8::GFP, UAS-mCD4::tdGFP, lexAop-mCD8::GFP and GMR-myr::tdTom (gift from Robin Hiesinger). *Trans-tango*: UAS-myr::GFP, QUAS-mtdTomato(3xHA)and trans-Tango ^13^
- longGMR-GFP::Hth ^43^ was used to transform retina to DRA

### Generation of transgenic DRA.R8-Gal4 and DRA.R8-LexA flies

The rh3 promoter containing three palindromic Prospero binding sites (Figure 1G; Dmitri Papatsenko, unpublished) was extracted EcoR1→BamH1 from a preexisting lacZ plasmid and ligated into promoterless Gal4-/ LexA-vectors with attP sites. Transgenes are inserted at attP2 (DRA.R8-Gal4) or attP40 (DRA.R8-LexA), respectively.

### Genotypes per Figure

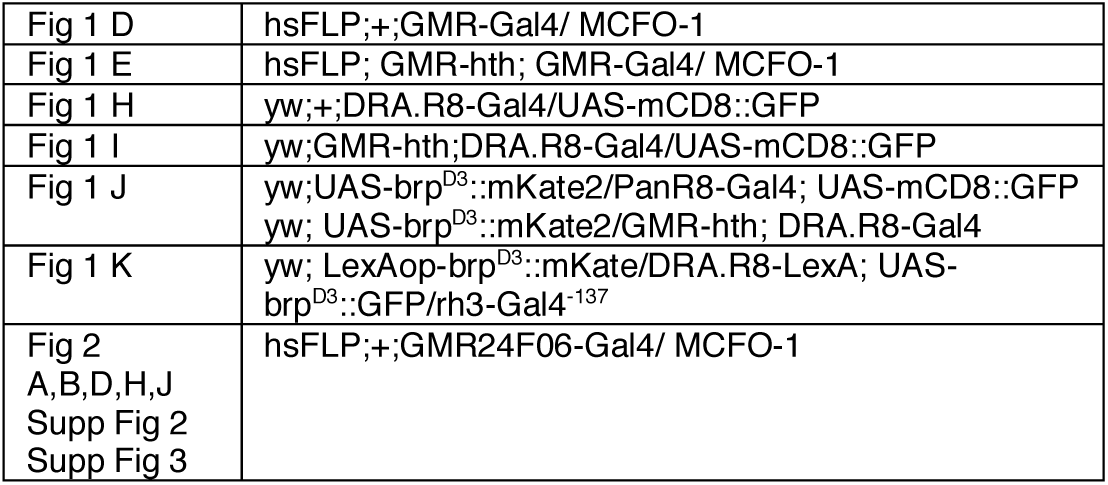
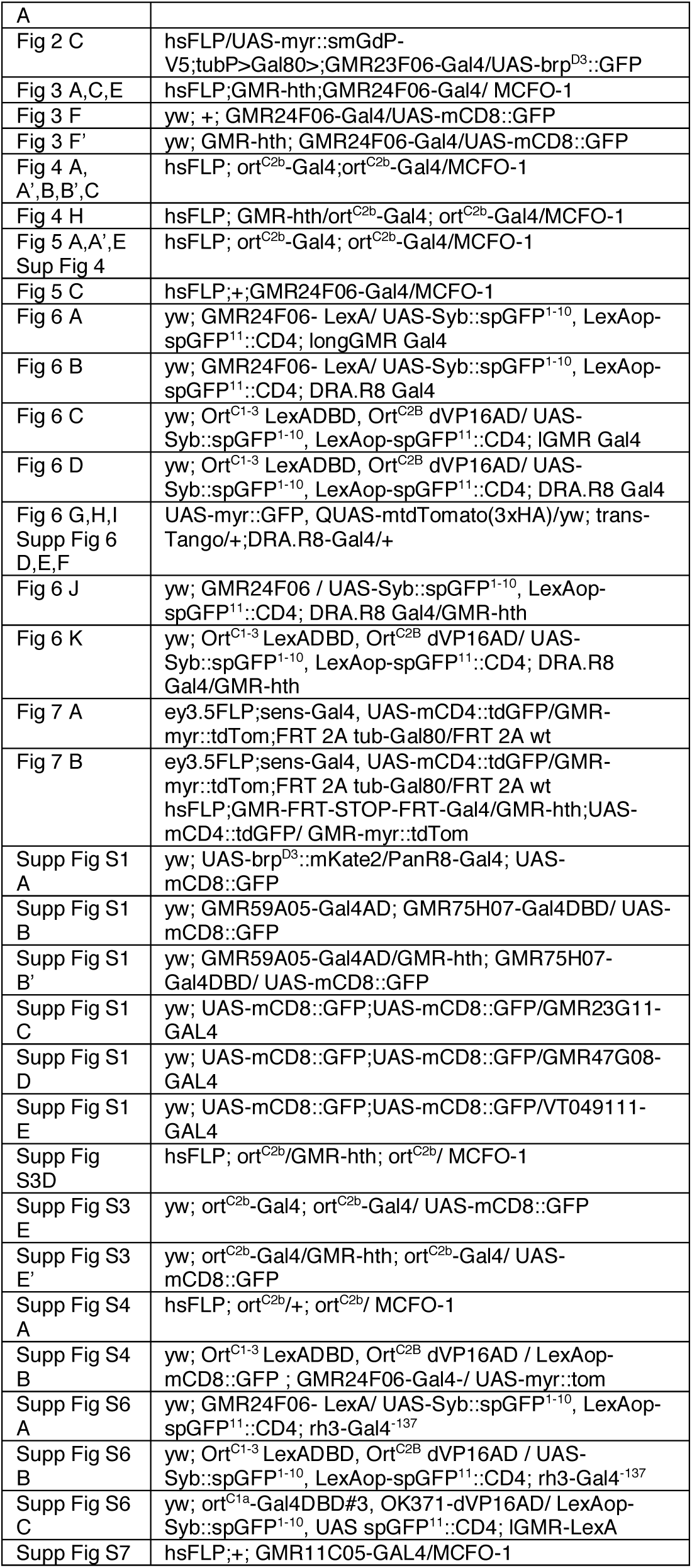

### Immunohistochemistry and imaging

Adult brains dissection was performed in ice-cold S2 cell culture medium (Schneider’s Insect Medium, Sigma Aldrich, #S0146) and brains were fixed with 4% PFA (v/w) in PBS for 20 - 30 minutes at room temperature. After 3 times washing with PBS-T [PBS with 0.5% (v/v) Triton X-100 (Sigma Aldrich, # X100)] fixed brains were incubated with primary antibody containing 10% Normal Donkey Serum in 0.4% PBS-T overnight at 4°C. Following three times washing with PBS-T, brains were incubated with secondary antibody solution containing 10% Normal Donkey Serum in 0.4% PBS-T overnight. After 3 times 15 minutes washing brains with PBS were mounted in Vectashield H-1000 (Vector Laboratory, Burlingame, CA) anti-fade mounting medium for confocal microscopy.

We used a Leica SP8 confocal microscope equipped with a white light laser and two HyD detectors. Image stacks were acquired in resolution of 1024×1024×0.5μm with a 63x lens.

### Antibodies used in this study

| Antibody | Host | Concentration | Company |
| --- | --- | --- | --- |
| 24B10 (Anti-Chaoptin) | Mouse | 1:250 | DSHB |
| Anti-HA | Rat | 1:250 | Roche |
| Anti-CD4 | Rabbit | 1:600 | Atlas Antibodies |
| Anti-dsRed | Rabbit | 1:500 | Takara |
| Anti-FLAG | Chicken | 1:1000 | Novus Biological |
| Anti-GFP mAb | Rat | 1:500 | BioLegend |
| Anti-GFP pAb | Rabbit | 1:1000 | Thermo Fisher Scientific |
| Anti-GFP pAb | Goat | 1:1000 | Abcam |
| Anti-HTH (dN-19) | Goat | 1:50 | Santa Cruz Biotechnology |
| DN-Ex #8 | Rat | 1:100 | DSHB |
| Rat-Elav-7E8A10 anti-elav | Rat | 1:50 | DSHB |
| V5 Epitope Tag Antibody Dylight™ 549 Conjugated | Rabbit | 1:1000 | Rockland |
| Anti-chicken Cy™5 | Donkey | 1:500 | Jackson Immuno Research |
| Anti-goat Alexa Fluor® 488 | Donkey | 1:500 | Jackson Immuno Research |
| Anti-goat Cy™5 | Donkey | 1:500 | Jackson Immuno Research |
| Anti-Rabbit Cy™3 | Donkey | 1:500 | Jackson Immuno Research |
| Anti-Mouse Alexa Fluor® 594 | Donkey | 1:500 | Jackson Immuno Research |
| Anti-Mouse Cy™5 | Donkey | 1:500 | Jackson Immuno Research |
| Anti-Rabbit Alexa Fluor® 488 | Donkey | 1:500 | Jackson Immuno Research |
| Anti-Rat Alexa Fluor® 488 | Donkey | 1:500 | Jackson Immuno Research |

### Single cell clones and MCFO

To obtain single cell clones and reveal the morphology and relative position of individual neurons in the adult visual system, tub-FRT-Stop-FRT with hsFLP and MCFO^11^ was used, respectively. 3 days-old flies were incubated in vials in a 37°C water bath for 30 minutes 3 days prior to fly brain dissection to induce flippase (FLP). To allow the expression of the reporter, the flies were kept over 3 days at 25° C. Dissection and staining occurred as described above.

### Activity GRASP

Flies were grown in a 25° C, 12h-12h dark-light cycle incubator in normal vials and transferred to custom-made UV-transparent Plexiglas tubes [wall thickness: 4mm, From RELI KUNSTSTOFFE (Kunststoffhersteller in Erkner, Brandenburg) (Gewerbegebiet zum Wasserwerk 16, 15537 Erkner)] before light induction. 1-day old flies were kept in a 25°C, 20 h – 4 h light-dark cycle custom-made light box (with UV light) for 3 days to ensure the photoreceptor activity in DRA.

Dissection & staining occurred as described above. Brains were stained with polyclonal GFP and monoclonal GFP antibody to visualize postsynaptic cells and GRASP signal, respectively. Post-synaptic cells were visualized by staining with CD4 antibody.

### Trans-Tango

Flies for trans-Tango experiment were either kept in 18° C or 25° C 12h light/dark cycle incubator and dissected when they were either 3, or 15 days old (depending on the experiment).

### Ex vivo live imaging

Ex vivo live imaging experiments were performed as previously described^42^. In brief, pupae were staged according to white pupae formation (0 APF) and dissected in ice-cold Schneider’s Insect Medium at the according time points. Pupal brain-eye complexes were cultured in a custom build culture chamber and perfused with oxygenated culture medium. Live imaging was performed with a Leica SP8 MP microscope with a 40X IRAPO water objective (NA = 1.1) with a Chameleon Ti:Sapphire laser (Coherent) at room temperature.

### Data Processing

#### Morphology and characteristics of the cells

Analysis and post-processing were done using IMARIS software. Single cell clones from MCFO data stacks were obtained by using the Surface and masking function in IMARIS. The cell location was determined by the angle (Supplemental Figure 2A). Contact points for Dm-DRA1 were determined according to photoreceptor surface and cell surface contact points and reviewed with fluorescent signal (Supplemental Movie 1). For Dm-DRA2 contact points were determined where photoreceptor surface and cell skeleton touch each other and reviewed with fluorescent signal (Supplemental Movie 2). Only one contact point determined per column. Length and width were determined with respect to contact point of each cell. Deep projections were determined as the projections below non-DRA R7 end (supplemental figure 2B). Vertical projections were the projections that contacts photoreceptors. Cell number analysis in GMR-hth experiment was done by manual counting of the cell bodies by Fiji.

#### Layer analysis

The row of photoreceptors belonging to the DRA was extracted by using the 3D Crop function in IMARIS. This row was then straightened by Fiji and a ROI that covered the end of R7 and R8 axons selected. To normalize the signal the image was binarized. The Average pixel intensity of the ROI was plotted for each channel (photoreceptor and cell).

#### Assembly of Dm-DRA clones

Skeletons from single cell clones were manually aligned to a common coordinate axis by using Track EM function of Fiji.

#### Live imaging

All 4D data stacks were deconvolved with the ImageJ (National Institute of Health) plug-in Microvolution (Microvolution, 2014-2016). The M0 layer, the corresponding R7 terminal and the deepest filopodium of individual R8 photoreceptor cell clones of each time point were manually marked with the spot function in IMARIS. The normalized reach into the medulla was plotted with R.

#### BRP distribution

Individual photoreceptor terminals were isolated with the surface, mask and 3D crop function in IMARIS. A new reference frame was placed into the approximate M0 layer of the isolated photoreceptor with the z-axis parallel to the photoreceptor terminal. BRP puncta were identified with the automatic spot function of IMARIS followed by manual revision. The distance of individual BRP puncta to the new reference frame was normalized to the deepest point of the R7 terminal and then plotted in R.

#### EM data analysis

EM data analysis (Figure 1L) was performed on two JSON data files downloaded from the Janelia webpage (http://emdata.janelia.org/#/repo/medulla7column). One containing all annotated synapses from the connectome and the other one all the annotated cell types. individual swc files were downloaded via the following webpage (http://emanalysis.janelia.org/gorgonian.php). A customized sharkviewer code was used to extract the synapse positions.

## Author Contributions

G.S., E.K., H.P. and M.F.W. designed the experiments and wrote the paper. J.B, L.M., T.P., A.H. and T.M. helped with experiments and data analysis.

## Acknowledgments

The authors are grateful to the late Dmitri Papatsenko who generated the original, previously unpublished DRA.R8 promoter sequence. The authors would like to thank Neset Ozel, Tiffany Cook, Maximilien Courgeon, Claude Desplan, Thomas Hummel, Abhishek Kulkarni, Marta Morey, Stephan Sigrist, and Aljoscha Nern for sharing fly stocks and reagents. Claude Desplan, Maximilien Courgeon, Tom Clandinin, Gerit Linneweber, and Robin Hiesinger provided helpful comments on the manuscript. This work was supported by the Deutsche Forschungsgemeinschaft through grants WE 5761/2-1 and SFB958 (Teilprojekt A23), through the Berlin Excellency Cluster NeuroCure, the US National Institute of Health (grant to Robin Hiesinger - division of Neurobiology, Freie Universität Berlin), the Fachbereich Biologie, Chemie & Pharmazie, as well as AFOSR grant FA9550-19-1-7005.

