## Supplemental Material for "Modality-specific circuits for skylight orientation in the fly visual system"

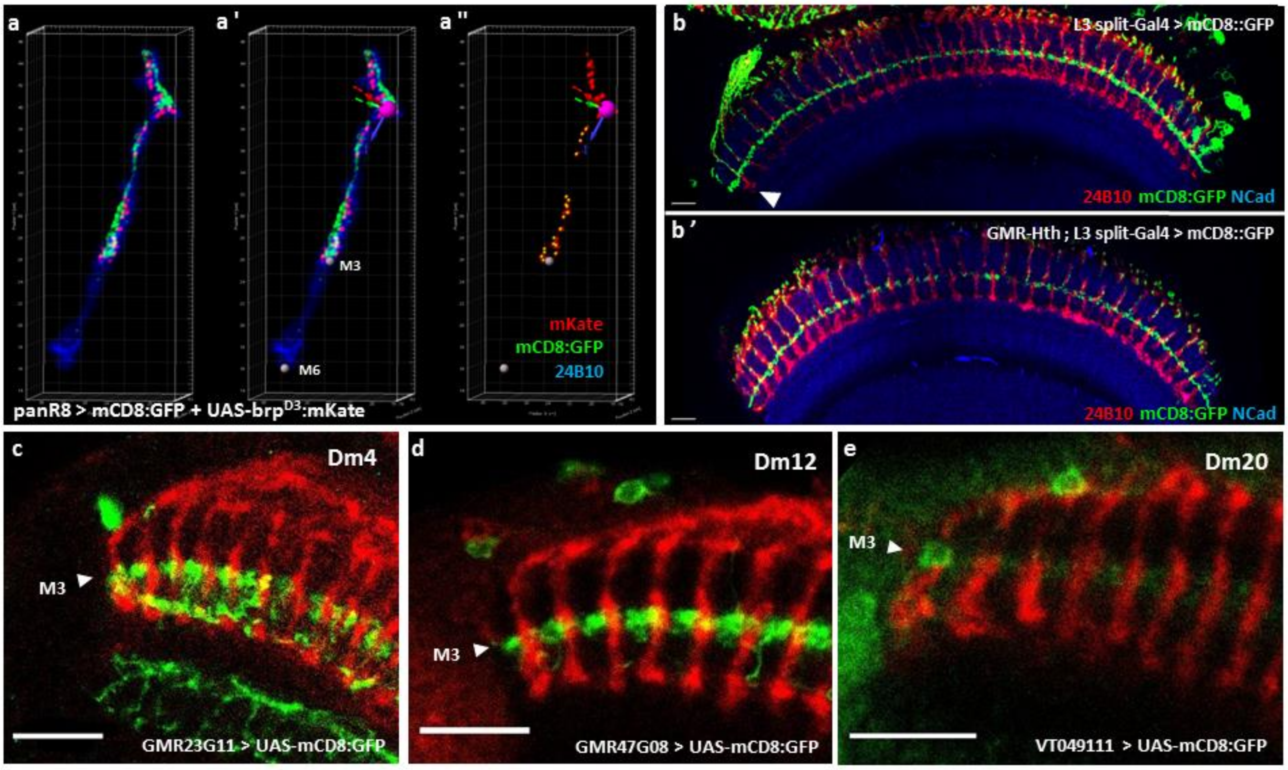


**Supplemental Figure S1: $Brp^{D3}$:XFP Distribution and medulla layers in the DRA. A**. Snapshots recapitulating the extraction of $Brp^{D3}$:mKate2 puncta using IMARIS. After extraction of signal for individual photoreceptors (A) a new reference frame was placed into the approximate M0 layer and two landmark points for M3 and M6 were added (silver balls) (A'). $Brp^{D3}$:mKate2 puncta were then identified with the automatic spot function (IMARIS) followed by manual revision (A''). **B.** layer targeting of lamina monopolar cell type L3 across the medulla: both inside the DRA region (arrowhead) as well as outside of it, L3 targets to the same layer (layer M3) in both wild-type (top) and GMR-hth background (bottom). **C-E.** Inside the DRA region, distal medulla cell types Dm4 (labeled with GMR23G11-Gal4) (C), Dm12 (labeled with GMR47G08-Gal4) (D), and Dm20 (labeled with VT049111-Gal4) (E) stratify normally within layer M3. Scale bars: 10 µm in (B and B') ; 5 µm in (C-E)

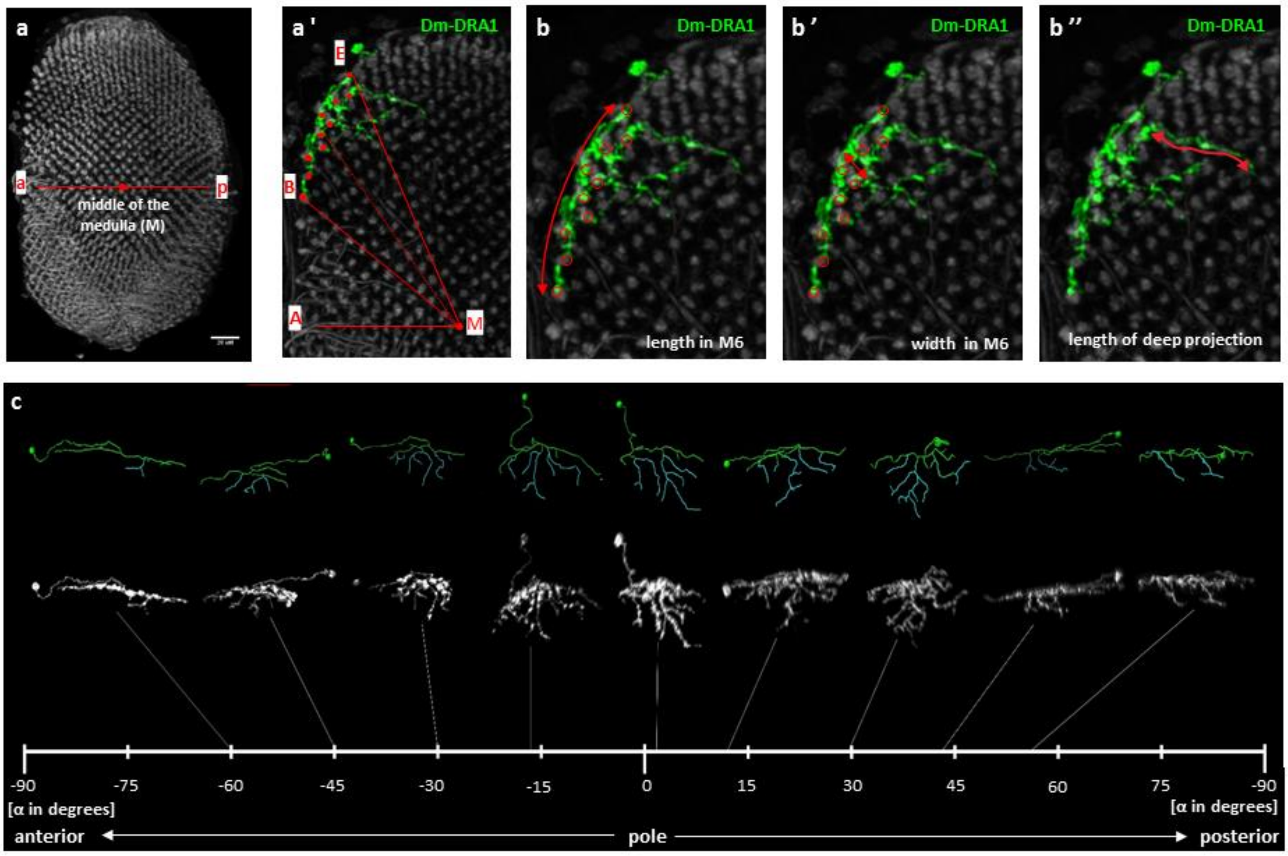


**Supplemental Figure S2: Morphometric criteria and characteristics of Dm-DRA1 cells. A**. Definition of middle of the medulla (M), from which the angular position α of a given cell along the DRA is measured. a: anterior, p: posterior. (A') Position of the cell calculated by $\widehat{AMB} + \frac{\widehat{BME}}{2}$ where B (beginning of the cell) is defined cell according to first photoreceptor contact point in M6 and E (end of the cell) is defined according to last contact point in M6; red balls show contact points. (Scale bar: 20 μm) **B.** Length of the arc in M6 spun by one Dm-DRA1 cell clone (B), reaching from points B to E, as determined from photoreceptor contacts, as above. Widest arborization of a given Dm-DRA1 cell, as defined by distance between photoreceptor cell contacts in M6 (B'). Length of deep projections (B'') is measured starting from the point where a process reaches below the photoreceptor terminals. Only the length of the longest process is quantified. Red balls show contact points. **C.** Gradual morphological change of Dm-DRA1 cell clones sorted from anterior DRA to posterior regions of the DRA. Green part of the skeletons marks the part of the cells with DRA photoreceptor contacts in M6. Deep projections are marked in cyan.

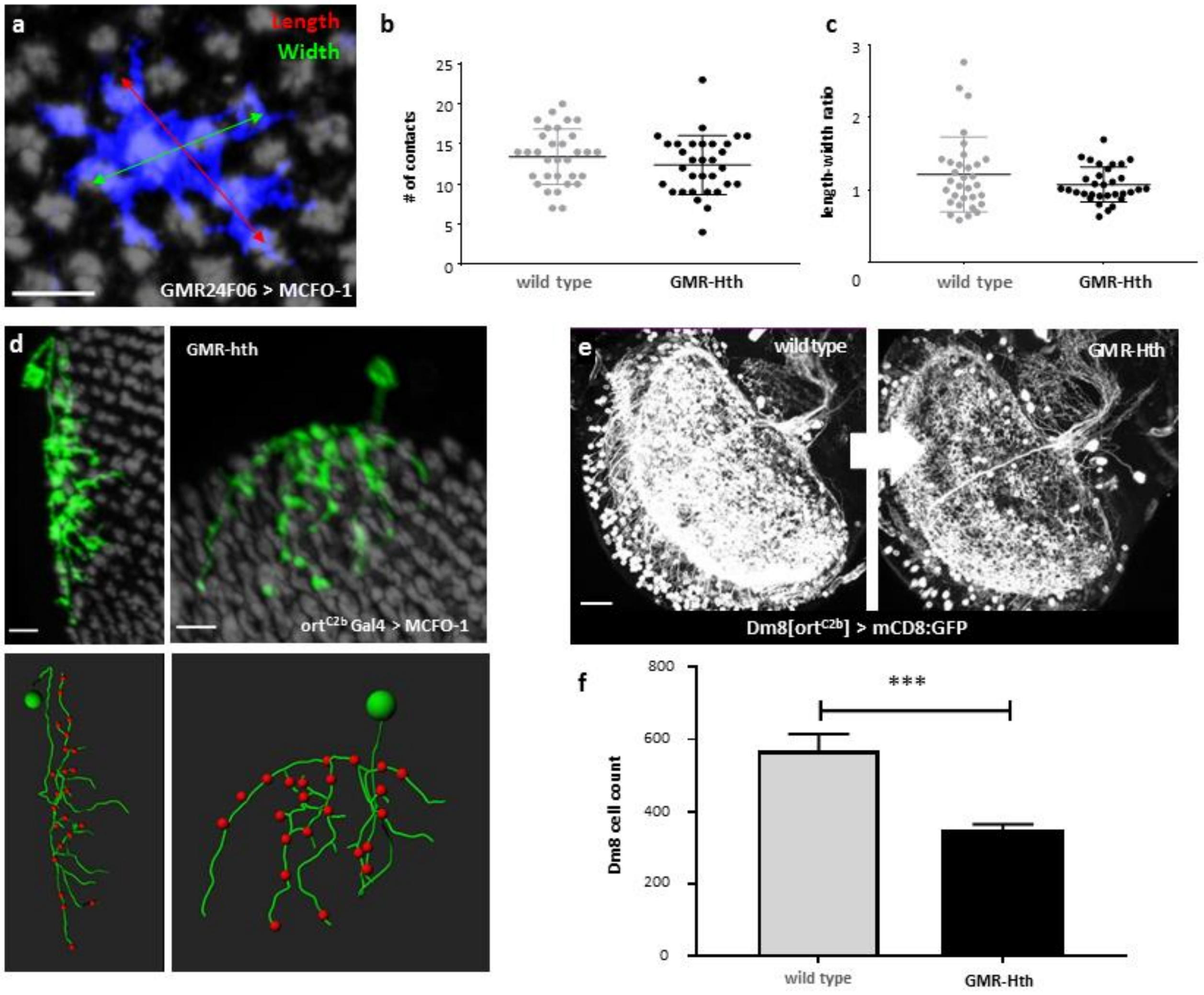


**Supplemental Figure S3: Effect of genetic redesign of the retina on non-DRA Dm8 cells. A.** The definition of length and width for non-DRA Dm8. Length is defined as the longest vector along dorsal-ventral axis of the cell where cell contact with photoreceptors. Width is defined as longest vector along anterior-posterior axis where the cells contacts photoreceptors. **B.** Quantification of photoreceptor contact number of non-DRA Dm8 in wild type and GMR-Hth flies. **C.** Comparison of cell shape as defined by the length/width ratio for non-DRA Dm8 in wild type and GMR-Hth flies. **D.** Top: Anterior (left) and polar (right) MCFO clones of Dm-DRA1 cells derived from ort$^{C2b}$-Gal4 driver. Bottom: Skeletons reconstructed from the above cells. Red balls show photoreceptor contact points in M6. **E.** Single-channel images of whole mounted ort$^{C2b}$-Gal4 > mCD8::GFP brains from wild type flies (left) and GMR-Hth flies (right), depicting the loss of Dm8 cells in the latter. **F.** Quantification of Dm8 cell number in wild type and GMR-Hth flies. Scale bars: 7 µm in (A and D) ; 20 µm in (E)

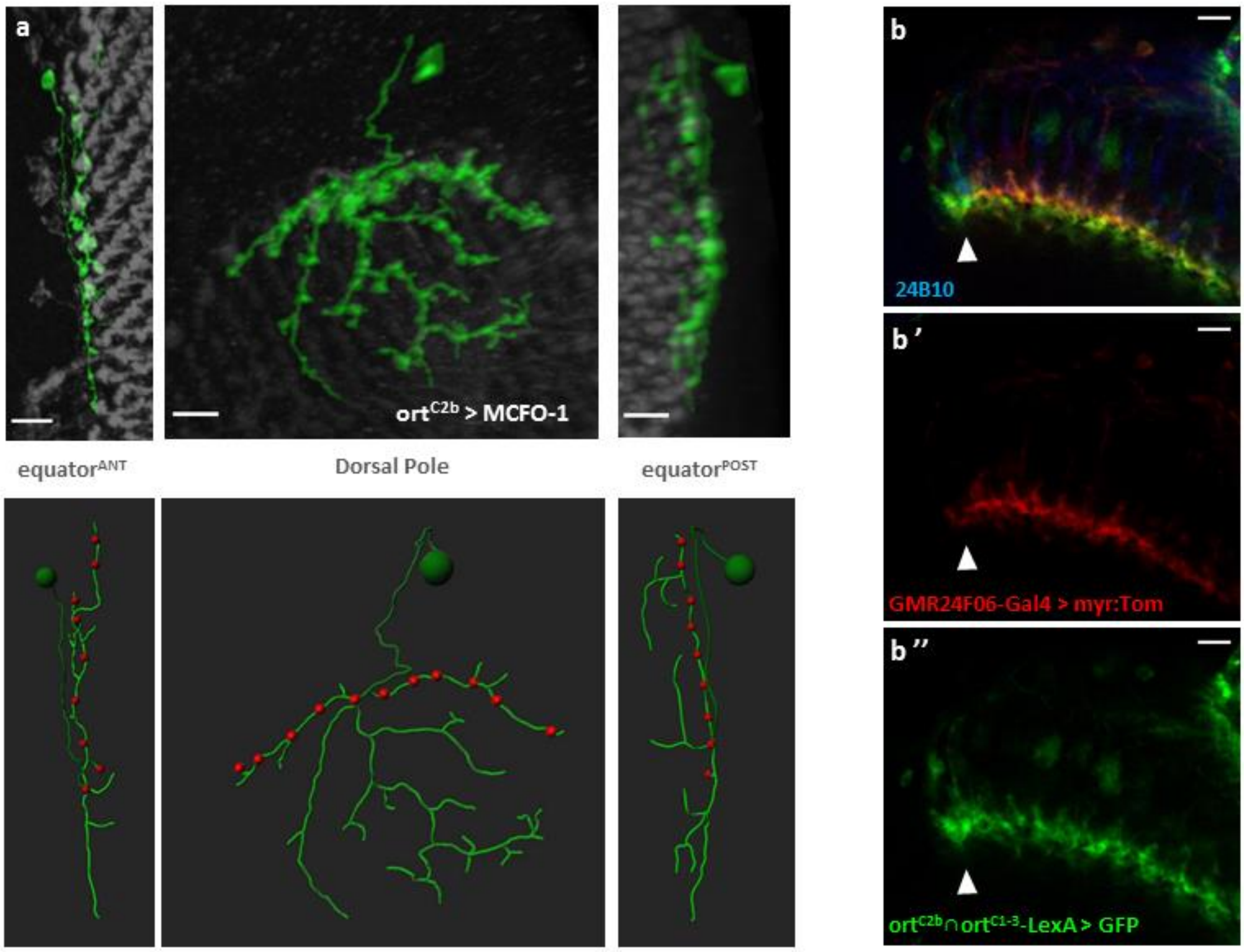


**Supplemental Figure S4: Expression of different Dm8 drivers in the DRA. A.** ort$^{C2b}$-Gal4 also labels Dm-DRA1 cells. Top: Three representative MCFO clones of Dm-DRA1 cells at different locations along the DRA: at the anterior equator (left), around the dorsal pole (middle), and at the posterior equator (right). Bottom: Skeletons reconstructed from the above cells. Photoreceptor contacts are visualized as red balls. **B.** Double labeling showing co-labeling of two Dm8 drivers (GMR24F06-Gal4 and ort$^{C2b}$∩ort$^{C1-3-}$LexA), resulting in differences in DRA columns (arrowhead). For comparison: single channel image of GMR24F06 > myr::Tom (B') lacks of vertical signal in DRA column which exist ort$^{C2b}$ ∩ort$^{C1-3}$ > mCD8::GFP (B''). Scale bars: 7 µm in (A and B)

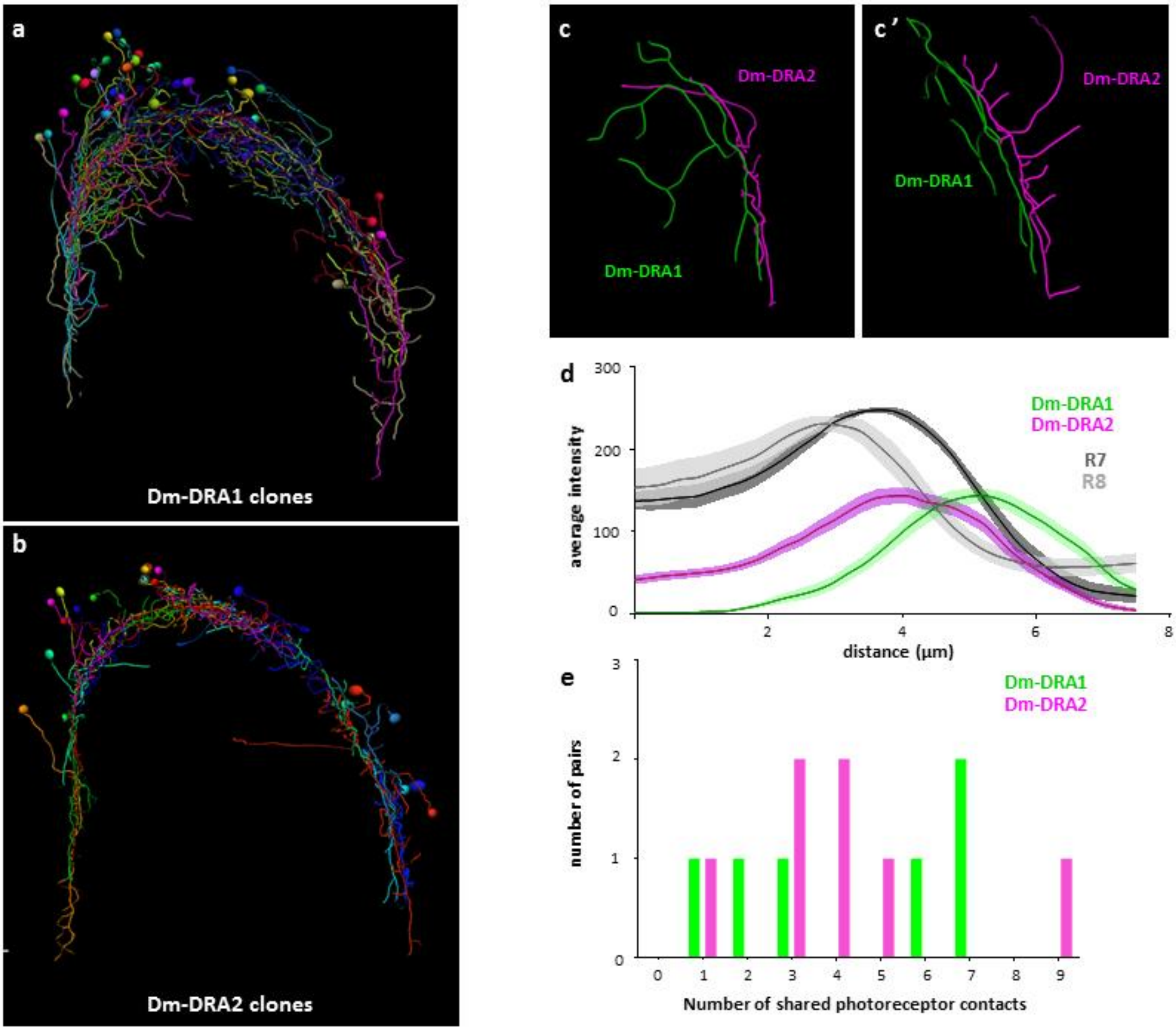


**Supplemental Figure S5: Both Dm-DRA cell types densely cover the DRA. A.** Artificially generated assembly of all Dm-DRA1 clones generated in different colors. **B.** Artificially generated assembly of all Dm-DRA2 clones in different colors. **C.** Skeletons of two adjacent Dm-DRA cell clones of different subtype (Dm-DRA1: green; Dm-DRA2: purple) located at roughly the same position in the DRA. Side view in C'. **D.** Quantification of R7 and R8 and Dm-DRA1 versus 2 layering, revealing positioning of Dm-DRA1 in close apposition but always distally from Dm-DRA2 similar to R7/R8 layering in the DRA, i.e. the distance between DRA.R7 and DRA.R8 (distal). **E.** Number of shared photoreceptor contacts between cell pairs of the same subtype, (Dm-DRA1: green, Dm-DRA2: purple).

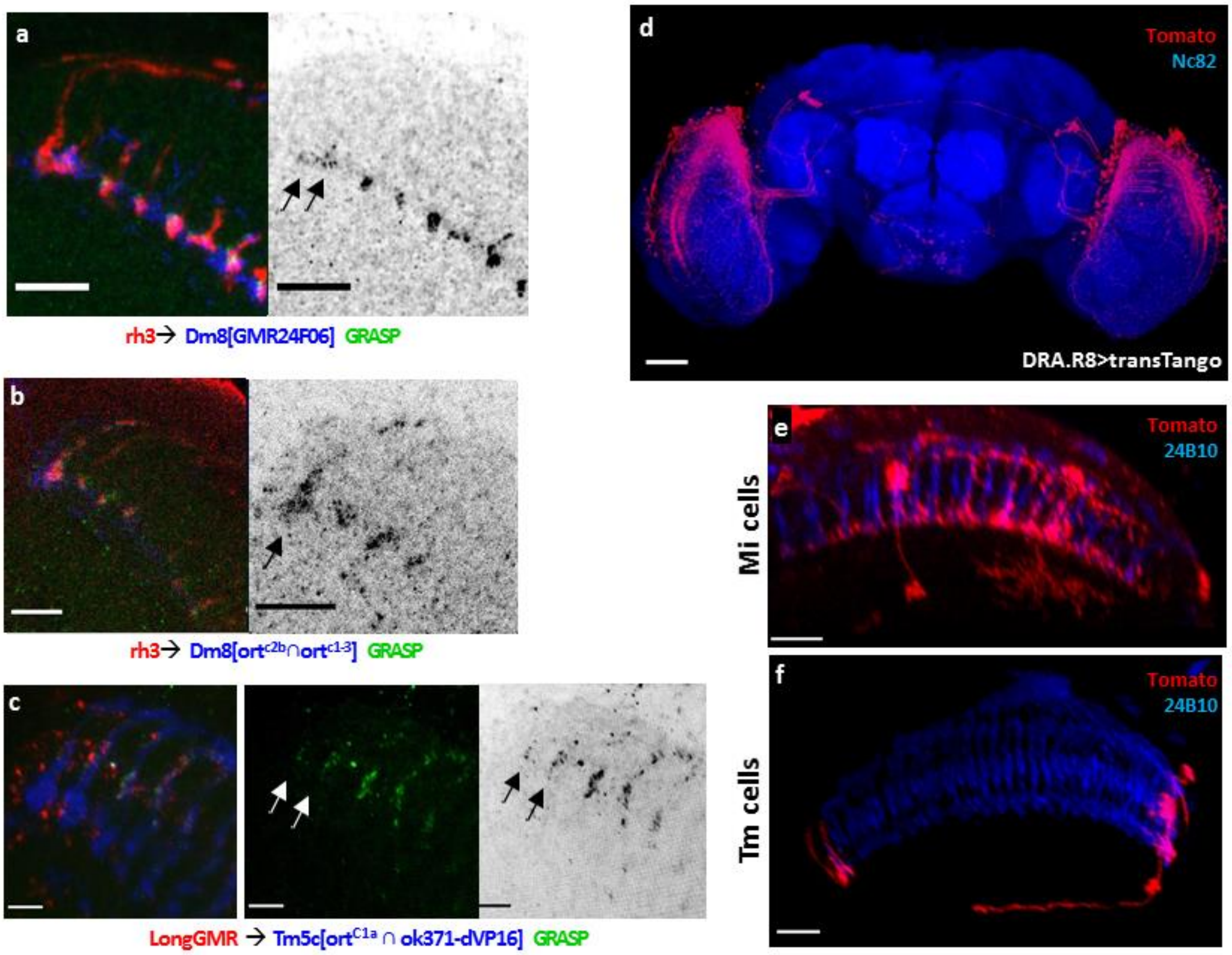


**<u>Supplemental Figure S6: GRASP and trans-Tango.</u> A.** Activity GRASP experiment visualizing potential synaptic contacts between photoreceptors (labeled with rh3-Gal4, UAS-nsyb:spGFP$^{1-10}$) and Dm-DRA1 cells (labeled with GMR24F06-LexA, LexAop-CD:spGFP$^{11}$). GRASP signal is restricted mostly to layer M6 throughout the medulla, including the DRA. **B.** Activity GRASP between photoreceptors (labeled with rh3-Gal4, UAS-nsyb:spGFP$^{1-10}$) and Dm-DRA1+2 cells (labeled with ort$^{c2b}$ ∩ ort$^{c1-3}$-LexA, LexAop-CD4:spGFP$^{11}$). GRASP signal spreads vertically in the DRA. **C.** Activity GRASP between photoreceptors (labeled with LongGMR-LexA, UAS-nsyb:spGFP$^{1-10}$) and Tm5c cells (labeled with ort$^{C1a}$ ∩ ok371-dVP16, UAS-CD4:spGFP$^{11}$). GRASP signal is present in the DRA, albeit weaker than outside of it. **D.** Saturated trans-Tango tracing (15 days, 18 degrees) using DRA.R8-Gal4 reveals many cells throughout the optic lobes. **E.** Example of Mi cell labeled by DRA.R8 > trans-Tango. **F.** Example of Tm cell labeled by DRA.R8 > trans-Tango. Scale bars: 10 µm in (A and B) ; 5 µm in (C); 50 µm in (D); 15 µm in (E and F)

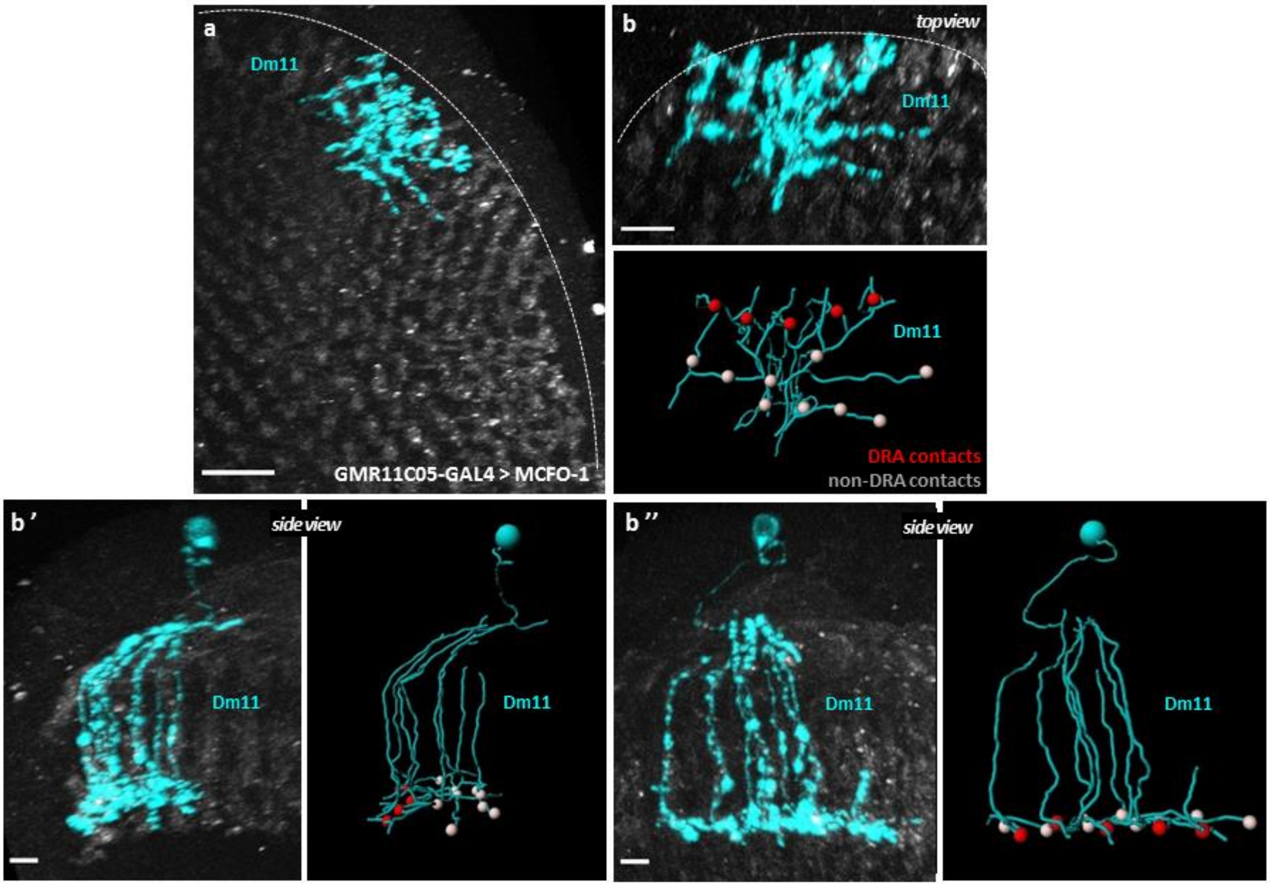


**Supplemental Figure S7: Morphology of Dm11 cells in the DRA. A.** MCFO clone of a Dm11 cell located at the dorsal edge of the medulla generated with GMR11C05-GAL4. **B.** Top view and side views (B,B',B'') of the Dm11 clone from A (cyan) with skeleton and photoreceptor contacts. Red balls show DRA photoreceptor contacts, silver balls depict contacts with non-DRA photoreceptor. Note that Dm11 cells do not respect the DRA boundary. Scale bars: 10 µm in (A); 5 µm in (B-B'')

### Supplemental movies

<u>Supplemental Movie 1: Photoreceptor contact points of Dm-DRA1 cells</u>

Animated movie describing how photoreceptor contact points were defined for the Dm-DRA1 cell type using IMARIS.

<u>Supplemental Movie 2: Photoreceptor contact points for Dm-DRA2 cells</u>

Animated movie describing how photoreceptor contact points were defined for the Dm-DRA2 cell type using IMARIS.

<u>Supplemental Movie 3: Live imaging of DRA.R8 and non-DRA R8 Photoreceptors</u>

16h live imaging of three R8 photoreceptor terminals in the dorsal edge of the medulla starting from P46 (individual photoreceptors were extracted with IMARIS, 2 frames/h.
